# Sedentary chromosomal integrons as biobanks of bacterial anti-phage defence systems

**DOI:** 10.1101/2024.07.02.601686

**Authors:** Baptiste Darracq, Eloi Littner, Manon Brunie, Julia Bos, Pierre-Alexandre Kaminski, Florence Depardieu, Weronika Slesak, Kevin Debatisse, Marie Touchon, Aude Bernheim, David Bikard, Frédérique Le Roux, Didier Mazel, Eduardo P.C. Rocha, Céline Loot

**Affiliations:** Institut Pasteur, Université Paris Cité, Unité Plasticité du Génome Bactérien, CNRS UMR3525, 75724 Paris, France; Sorbonne Université, Collège doctoral, F-75005, Paris, France; Institut Pasteur, Université Paris Cité, Microbial Evolutionary Genomics, CNRS UMR3525, 75724 Paris, France; DGA CBRN Defence, 91710 Vert-le-Petit, France; Institut Pasteur, Université Paris Cité, Synthetic Biology, CNRS UMR6047, 75724 Paris, France; Institut Pasteur, Université Paris Cité, Molecular Diversity of Microbes lab, INSERM U1284, 75724 Paris, France; University of Montréal, Faculty of medicine, Department of microbiology, infectiology and immunology, Montreal, QC H3T 1J4, Canada

**Keywords:** genetic adaptation, bacterial immunity, streamlined anti-phage system, *Vibrio* spp

## Abstract

Integrons are genetic systems that accelerate bacterial adaptation by acquiring and shuffling gene cassettes. Mobile integrons spread antibiotic resistance genes among bacteria, while the sedentary chromosomal integrons contain up to hundreds of cassettes of unknown function. Here, we show that many of these cassettes encode anti-phage defence systems. We found numerous streamlined variants of known systems, which have presumably evolved to fit the small size constraints of integron cassettes’ recombination and genesis. Intrigued by the rarity of known systems in the sedentary chromosomal integron of the *Vibrio cholerae* 7^th^ cholera pandemic strain, we tested the presence of anti-phage functions in all its cassettes of unknown function. We found that at least 16 of the strain cassettes have an anti-phage activity in *V. cholerae* or *E. coli*. This represents 18% of the tested cassettes and almost 10% of all the integron cassettes, providing at long last a key adaptive role for a significant fraction of the sedentary integrons. Most of the newly discovered systems have little or no similarity to previously known ones and our experiments show that several mediate defence through cell lysis or growth arrest. One of these systems encodes a 64 amino acids protein, which represents the smallest known protein providing autonomous phage resistance. Given the thousands of uncharacterized integron cassette families, integrons could represent an untapped treasure trove of streamlined anti-phage systems.

## INTRODUCTION

Unexpected challenges require fast responses that may be difficult to prepare. This is especially true for antagonistic biotic interactions, such as the attacks bacteria endure by bacteriophages (phages) (*1*). Many anti-phage systems have evolved to cope with this challenge (*2*). They constitute a fast-changing pangenome of defence systems extensively exchanged within bacterial communities by horizontal transfer (*3–5*). The high turnover of these functions is caused by the huge genetic diversity of phages and other MGEs, and by their ability to evolve anti-defence systems (*6*). Defence systems are under positive selection when targeting co-occurring infectious phages but may be counter-selected when their costs are not sufficiently compensated by their protective role. The latter tends to fade away as phages evolve to become resistant, which is thought to result in cycles of gain and loss of defence functions. Yet, gene gain by horizontal transfer evolves in a much longer time scale than that of phage infections. If a bacterial genetic system could integrate and express adaptive functions at a given moment while storing them at low cost when not needed, then bacteria could rapidly respond to phage pressure without relying on hazardous horizontal gene transfer. We postulated that integrons could perform these multiple roles: facilitate acquisition defence systems, archive them as a biobank and orchestrate their stochastic expression or silencing. They could thus facilitate bacterial’s response to the challenges posed by phages, which will inevitably infect bacteria, but whose diversity and rapid evolution complicate the evolution of a fixed set of defence systems. Integrons are genetic structures that capture, stockpile, shuffle and express promoter-less gene cassettes terminated by recombination sites (*attC*) (*7*). These events are mediated by a peculiar recombinase, the IntI integrase that acts in two ways: it excises cassettes from the array, and it integrates them at its beginning (*attI* site) where they become expressed under the action of the integron promoter (P_C_) (Figure 1A). Cassette integrations gradually shift the other cassettes away from the promoter. The cassettes may thus remain silent and costless for large periods of time until they are mobilized to the expression site (*7–10*). Recombination between *attC* sites decreases as the length of the cassettes increases (*11*). This is thought to constrain the functions of genes present in integron cassettes, since the latter are significantly smaller than the average gene (*11*). The integron integrase is expressed under stress conditions allowing combinatorial sampling of the phenotypic diversity (*12*). The configurations of the array of cassettes that increase bacterial fitness in the local conditions will then be selected and rise in frequency in the population. This stochastic system of adaptation on demand could fit exquisitely the bacterial requirement for a reservoir of anti-phage systems. Integrons are commonly divided in two types (Figure 1B). Mobile integrons (MIs) have typically less than a dozen cassettes, often encoding antibiotic resistance genes, and are associated with MGEs (*13*). Sedentary chromosomal integrons (SCIs) are not mobile. Their presence is conserved across bacterial species, even though their repertoire of cassettes may change rapidly. SCIs encode large arrays of gene cassettes (up to more than 200), most of which of unknown function (*14, 15*). Yet, it was previously shown that integrons over-represent genes matching the COG categories including toxin-antitoxin, restriction-modification, and other defence systems (*14–16*). The toxin-antitoxin systems were experimentally shown to stabilize the integron by their addictive behavior (*17*), but they could also be anti-phage systems (*18*).

**FIGURE 1:**
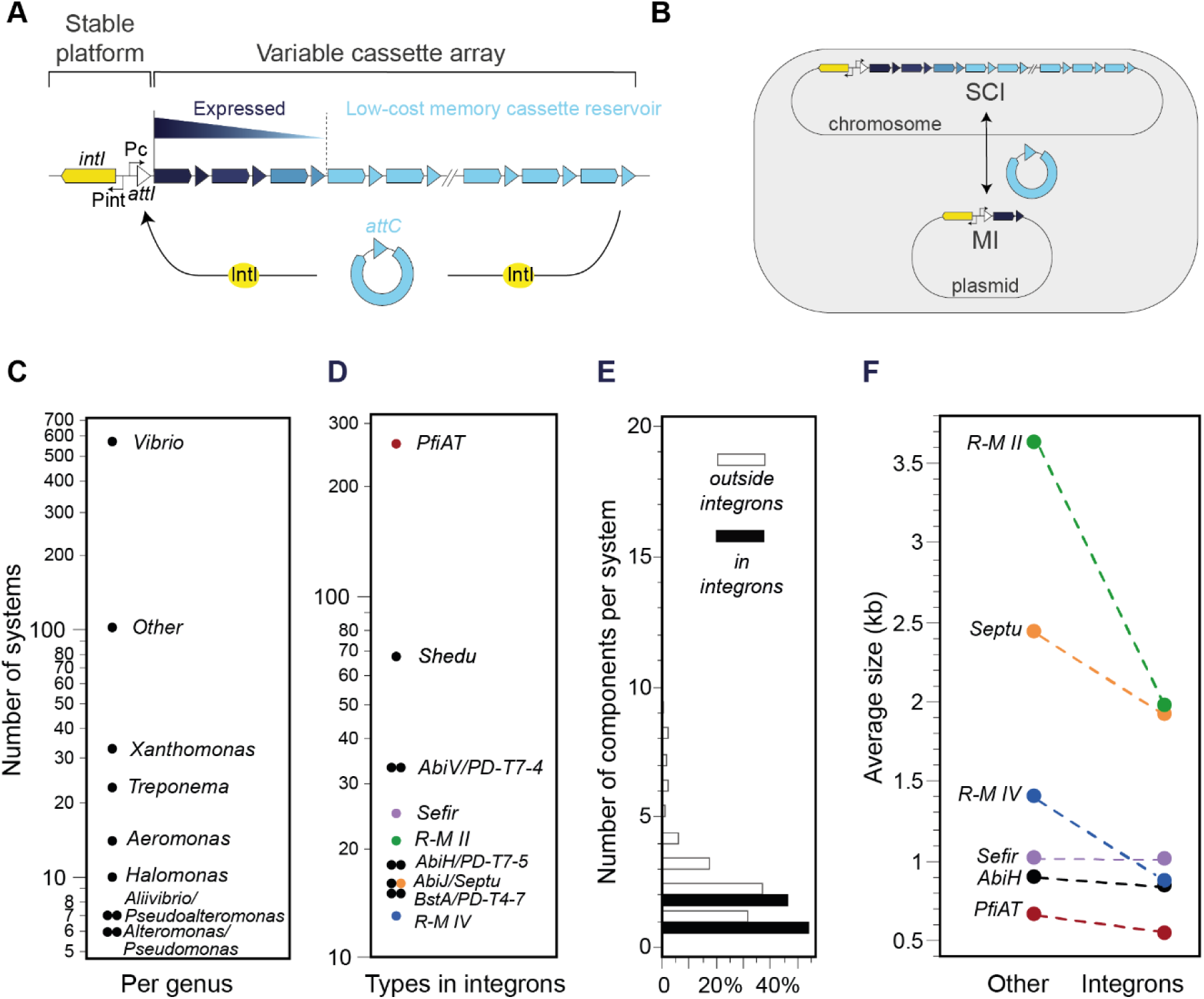
Integrons and their known anti-phage systems. **A.** Representation of the integron system and function. The stable platform consists of a gene encoding the integrase (*intI*, yellow) and its P_int_ promoter, the *attI* cassette insertion site (white) and the P_C_ cassette promoter which drives the expression of downstream cassettes along an expression gradient. The cassettes are represented by blue arrows and triangles: arrows represent the CDS and triangles, the *attC* sites. The intensity of their color means their level of expression, from dark blue (highly expressed) to light blue (not expressed). Cassettes can be excised by an intramolecular *attC* × *attC* reaction and reintegrated into the 1^st^ position near the P_C_ promoter by an intermolecular *attC* × *attI* reaction and become expressed. **B.** This process may also result in cassette exchange between Sedentary chromosomal and mobile integrons (SCIs and MIs). **C-D.** Number of systems identified in integron cassettes in function of the taxonomy and in function of the type of system. **E.** Number of protein coding genes (components) identified per anti-phage system when comparing systems identified in integrons with systems identified in other regions of the bacterial genomes (all genomes of the database). **F.** Average size of anti-phage systems identified in replicons with integrons, separating the systems encoded in integrons from the others. To address taxonomic biases, we compared systems within similar genetic backgrounds. Thus, for the analysis displayed in this panel, we only included systems present in replicons with at least one integron-located defence system. Only system types with more than 10 occurrences in integrons and more than 10 occurrences outside integrons are represented.

Here, we explore the idea that integrons are biobanks of anti-phage systems. We start by identifying many known or putative defence systems in integrons, especially in the SCIs. Intriguingly, only one was identified in the large (179 cassettes, 130 kb) model SCI of the seventh pandemic *Vibrio cholerae* O1 El Tor strain, N16961. We therefore hypothesized that this SCI might harbor unknown anti-phage functions. We independently cloned all non-redundant cassettes of unknown function and challenged them against phages of *V. cholerae* and *E. coli*. We found that many of these cassettes provide resistance to phages. We performed computational annotations on these cassettes and subsequently conducted experiments demonstrating that several of them induce cell lysis or growth arrest, preventing phage amplification. We postulated that the significance of integrons hinges on their capability to position cassettes downstream of an active promoter and validated that relocating cassettes to the first position in the array indeed confers protection. Our results confirm that integrons are cost-effective biobanks and orchestrators of anti-bacteriophage defences, highlighting their potential as untapped treasure troves of defence systems.

## RESULTS

### Sedentary chromosomal integrons have numerous streamlined anti-phages systems

We searched the complete reference genome database RefSeq (Table S1) for known or putative defence systems using DefenseFinder (*19*) and identified 218,635 systems in 32,798 genomes (Table S2). In parallel, we used IntegronFinder on the same dataset and identified 85,204 integron cassettes comprising 121,711 protein-coding genes (Table S3). The intersection of both analyses revealed that 783 complete defence system are encoded within integron cassettes. The frequency of defence systems in integrons was 2.2 times higher than expected given the distribution of these systems in genomes (P<0.001, χ^2^ test). The defence systems in integrons show a peculiar taxonomic distribution, since the vast majority were found in *Vibrio* (73%, Figure 1C). This is largely because this clade has the largest number of cassettes (60%, Figure S1). One could have thought that such systems were present in MIs which are associated with MGEs. Instead, SCIs, representing 86% of the cassettes, account for over 99% of the identified defence systems. This may be the consequence of the over-representation of MIs carrying antibiotic resistance in current databases (leaving the other MIs under-represented). We conclude that recognizable defence systems are abundant in integrons, especially in the large chromosomal ones.

We then investigated the types of defence systems identified in integron cassettes (Figure 1D, Table S4). By far the most common systems are toxin-antitoxin (TA) systems (a third of all systems) of the PfiAT family (*20*). TAs also have properties of poison-antidote systems making them addictive and thereby capable of stabilizing the repertoire of cassettes of SCIs (*17*). The SCI of *Vibrio cholerae* El Tor str N16961 - belonging to the current 7^th^ pandemic lineage and chosen as the focal strain of this study, see below - harbors 19 TAs (*21*). Among them, the relB/parE-like TA (VCA0488/0489) was detected as a PfiAT defence system. More generally, all the SCIs carried by 7^th^ pandemics *V. cholerae* harbor a PfiAT and no other defence systems (Figure S1). The second most frequent defence systems are Shedu systems which are proposed to act as nucleases (*2, 22*), followed by an abortive infection system (AbiV) and by PD-T7-4 whose function is yet poorly understood.

Interestingly, the genetic loci encoding these systems are small, TA systems are usually composed of two short genes and the remaining systems are composed of a single gene. Since the integron cassettes contain smaller CDSs than replicons probably due to the cassette genesis and recombination constraints (*11*), we examined the hypothesis that the pool of defence systems in integrons comprises systems with a smaller-than-average number of components. Indeed, the types of systems encoded in integrons tend to be smaller than the average system identified across the genome database and never exceeds 2 components (Figure 1E). We then enquired if across the types of systems identified in integrons, they tended to be smaller than the same types of systems found at other locations in the same genomes. For this, we built a mixed model describing the variation in the length of the system where the presence in or outside the integron was the fixed effect and the type of system was the random effect. The results show that length is significantly smaller for systems in integrons when controlling in this way for the effect of the type of system (R^2^=0.87, P<0.0001, F Test, Table S5 and S6). This trend becomes particularly evident when one compares the average size of systems present in more than 10 occurrences both within and outside integrons (Figure 1F). Hence, these results show that the defences encoded in cassettes are smaller than expected, both when looking at systems (reduced number of components) and at featured genes themselves (shorter CDS). The need for gene compaction to facilitate recombination within the integron may explain why smaller restriction systems (such as type II and IV) are more prevalent, while defence systems featuring lengthy and intricate loci encoding anti-phage mechanisms like CRISPR-Cas or BREX are entirely absent from integrons.

### Discovery of unknown anti-phage systems in the *Vibrio cholerae* SCI

The previous analyses revealed few defence systems in the integrons of the 7^th^ pandemic strains, which we found intriguing. This led us to investigate the anti-phage activity of the 88 non-redundant function-unknown cassettes of the SCI carried by the *V. cholerae* El Tor str N16961, the model SCI for *V. cholerae*’s 7^th^ pandemic. Since these cassettes are poorly or not expressed under our laboratory conditions (*23*), we cloned them under an inducible promoter in a plasmid (Methods). Each construct was transferred to a *V. cholerae* derivative lacking both the SCI (to avoid interference with other cassettes) (*24*) and the anti-phage system DdmABC (to recover the phage X29 activity, Figure S2) (*25*). We then challenged the bacteria with our collection of five phages: ICP1, ICP2, ICP3, 24 and X29. We observed a reduction in plaque number and/or a marked reduction in plaque size, both phenotypes indicative of altered susceptibility to phages. Eight cassettes provided robust and reproducible resistance to one phage. One cassette alters the susceptibility to three phages out of five (Figure 2A, Figure S3).

**FIGURE 2:**
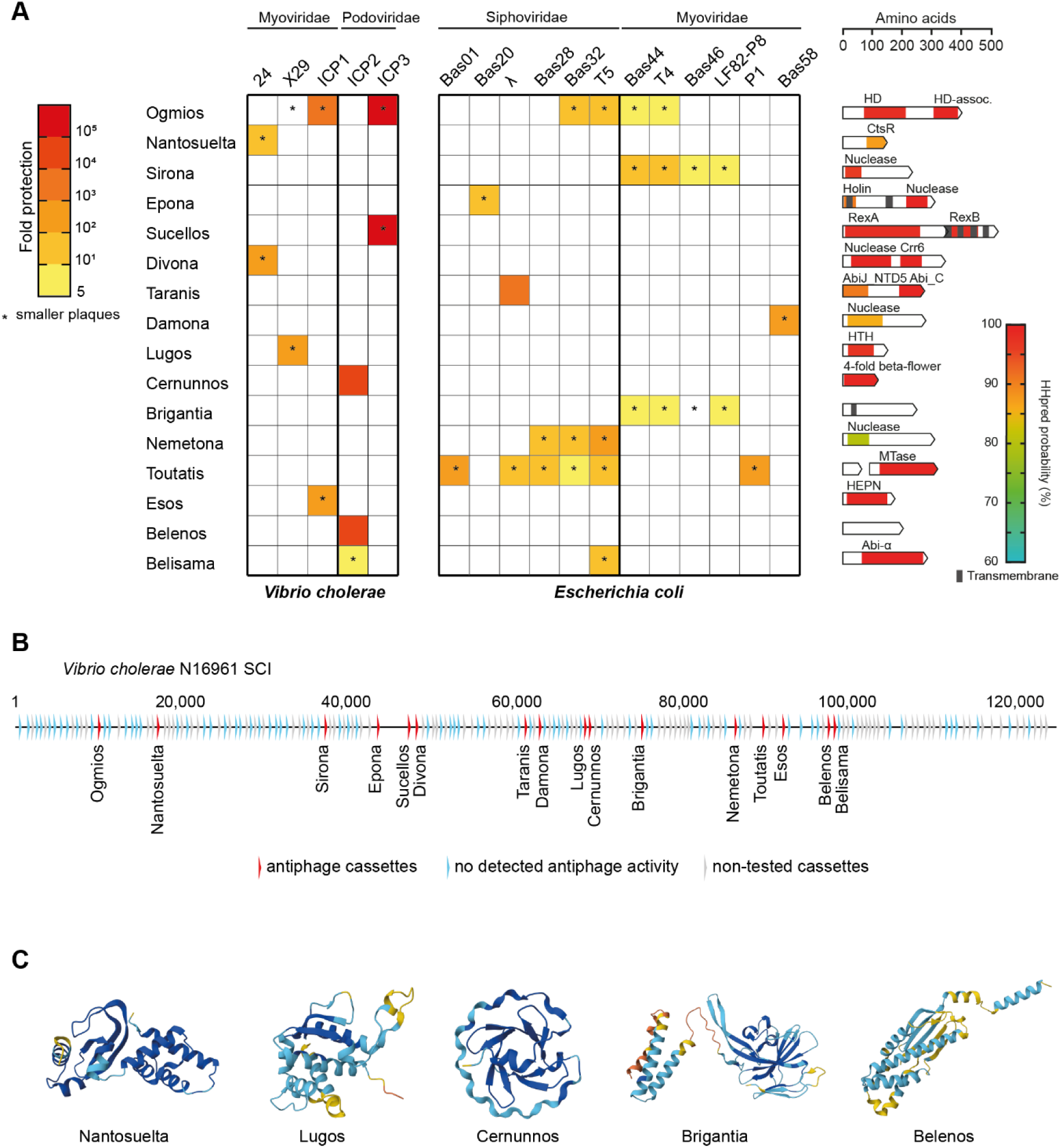
*V. cholerae* SCI cassettes protect against phages. **A.** Heatmap of the phage resistance phenotype representing the mean fold resistance of three independent replicates against a panel of 17 *V. cholerae* or *E. coli* phages. Systems are represented in lines and homology to known protein domains is displayed on the right. Phages are represented in columns and classed according to morphology. In two cases – Sucellos and Toutatis – two genes are in the same integron cassette. * stands for a marked reduction in plaque size. **B.** Position of the identified anti-phage cassettes in the *V. cholerae* SCI (red), those that were tested but did not reveal activity (blue), and those not tested (grey). **C.** Predicted structures (AlphaFold2) of five anti-phage cassettes in the *V. cholerae* SCI. The predicted structures of all anti-phage cassettes are depicted in Figure S4.

The exploration of the anti-phage function of 88 cassettes remains limited in *V. cholerae* due to the scarcity of available phages and their potential previous adaptation to the strain. It is also useful to note that integron cassettes are commonly transferred across bacterial species, via MIs, and could confer anti-phage protection to a wide range of hosts. Consequently, we sought to re-evaluate cassette function using a “naive” heterologous system, *E. coli* MG1655, for which a much broader panel of phages is available. Each cassette was expressed in *E. coli* MG1655 and challenged with a collection of 23 phages. Five cassettes offer protection against 3 to 5 phages, whereas 4 cassettes seem to be more phage-specific (Figure 2A, Figure S3). Anti-phage activities were observed in both hosts for two cassettes. We named these 16 verified unknown anti-phage systems after deities from Celtic mythologies (Figure S4). Overall, 16 out of the 88 cassettes (18%), representing 10% of all integron cassettes, worked as anti-phage systems in either the *E. coli* or the *V. cholerae* strains (Figure 2B).

### Description of the anti-phage systems found in the *Vibrio cholerae* SCI

None of the 16 anti-phage systems that we identified in *Vibrio cholerae* SCI cassettes was detected by DefenseFinder using default parameters. Defence systems are often a modular combination of domains (*e.g.* a sensor and an effector) (*26–28*). We thus investigated whether the newly identified systems had protein domains found in known systems using available data (Uniprot, (*29*)), sensitive sequence similarity detection programs (HHPred (*30*)), and structural similarity (AlphaFold (*31*) and FoldSeek (*32*)) (Figure 2C, Figure S4). Of note, as expected given our observation above that known defence systems in integrons tend to be small, all the discovered systems are small: two have two genes, all others have a single one.

Five systems have domains also found in systems previously described causing growth arrest or cell death upon infection (*33*). Ogmios encodes a putative deoxyguanosine triphosphatase (dGTPase) enzyme, which degrades dGTP into phosphate-free deoxyguanosine. A dGTPase was previously shown to deplete the nucleotide pool available for phage assembly (*34*). A homolog of Ogmios, classified as antiviral by DefenseFinder, was actually detected elsewhere in the genome of *V. cholerae* El Tor str N16961. Ogmios may hence be part of a larger family of antiviral dGTPases.

Sucellos has high predicted structural similarity to RexAB encoded by phage *λ* (*35*), even though it is not matched by the respective PFAM domain. RexA senses phage infection and triggers the membrane-anchored protein RexB, an ion channel which destroys the membrane potential resulting in cell death (*36*). Since the RexAB function requires the coordinated action of RexA and RexB, we tested separately the anti-phage function of the two genes of Sucellos (that we named *sclA* and *sclB*) and, as expected, found that both are necessary to provide anti-phage activity (Figure S5A and B).

Taranis may have a function related to that of AbiJ since it matches the PFAM domain Abi_C, corresponding to the C-terminus of the *Lactococcus lactis* abortive infection protein Abi-859 (*37*), a system later renamed AbiJ (*38*). The C-terminus of AbiJ is conserved and belongs to the superfamily of HEPN domains that is common in defence systems (*39*), especially those involved in growth arrest or cell death upon infection (*40*). Conversely, its N-terminus can vary. The N-terminus of Taranis is distantly related to AbiJ_NTD5. Several systems containing an Abi_C or an AbiJ_NTD domain were recently identified in phage-inducible chromosomal islands (PICIs) of *Klebsiella* (*41*).

Esos has only a recognizable HEPN-like domain and no other detectable feature. Finally, Belisama has a DUF2806 that has detectable homology with an Abi-α domain, an abortive system often found in prophages (*42*).

Five anti-phage defence systems harbor diverse fusions of nuclease domains that might act as effectors and other protein domains. Epona harbors a catalytic domain (Mrr_cat) found in type II and type IV restriction enzymes (*43*), fused to a holin domain with 2 transmembrane domains. Of note, an Mrr_cat domain can also be found in other recently described defence systems’ effectors such as Lamassu, Avs and NLR-related systems (*44–46*). Divona has an unknown function domain (DUF4365) in the N-terminus and a RecB-like nuclease of the Mrr subfamily (*47*), fused to a chlororespiratory reduction 6 (crr6, PF08847) domain. To our knowledge, the latter was not previously found in anti-phage defence. Damona harbors a domain DUF6602 which is distantly related to RmuC. The latter is a key domain involved in the defence system Shield (*48*), where it has non-specific (endo)nuclease activity for phage, plasmid, and chromosomal DNA. Damona is further matched by two domains involved in more complex defence systems relying on nuclease activity: CRISPR-Cas (Cas PD_DExK) and CBASS (CBASS_Endonuc_small). Sirona and Nemetona are both distantly related with His-Me finger endonucleases (HNH), notably to a type II restriction endonuclease domain and to phage *λ* NinG protein (PF05766). NinG (or Rap) increases the *λ* phage recombination catalysed by *Escherichia coli*’s RecBCD pathway (*49*).

Toutatis has two genes, *tutA* and *tutB*, but was considered as a single system because the two genes co-localize in the same cassette in all 61 identified occurrences in *V. cholerae* genomes. *tutB* is a putative methyltransferase. *tutA* is homologous to one domain of the restriction endonuclease Hpy99I involved in cleavage site recognition (*50*), but does not contain catalytic residues. We tested separately the anti-phage function of the two genes and found that *tutA* is necessary and sufficient to provide anti-phage activity (Figure S5C and D). To our knowledge, the small protein (64 amino acids) encoded by *tutA* is the smallest known to provide autonomous phage resistance.

Five systems have no homology to domains previously associated with defence systems (Figure 2C). Nantosuelta is distantly related to the C-terminal dimerization domain of CtsR (PF17727), a protein that controls heat shock gene expression in *Listeria monocytogenes* (*51*). Lugos is distantly related to helix-turn-helix DNA-binding proteins. Cernunnos harbors a 4-fold beta flower domain (PF21784) recently described (*52*). Brigantia has a transmembrane domain and does not match any other domain of known function. Finally, Belenos lacks known functional domains and homologs of known function. In summary, 11 systems have protein domains or homologous protein domains that can be found in known defence systems, whereas 5 are completely unknown. Their molecular mechanisms will have to be determined in the future and we are currently concentrating our attention on Toutatis system.

### *V. cholerae* anti-phage systems trigger cell lysis or growth arrest

We investigated the function of the systems that are effective in their natural *V. cholerae* host. We monitored growth of cells carrying each of the nine systems or a control plasmid during infection at a low (0.01) or high (10) MOI. Bacterial populations resisted well to phages at low MOI (Figure 3A). Interestingly, at a high MOI, all but one (Nantosuelta) display cell lysis or halted growth after infection. While Lugos and Belisama elicit a strong and unrecoverable culture lysis, most of the others reveal short growth arrest followed by regrowth, which could be associated to post-infection dormancy (Figure 3A) (*33*). Phage quantifications after lysis when infected at high MOI show an absence of amplification (Figure 3B). Therefore, all systems tested here, besides Nantosuelta, seem to halt growth or kill infected cells preventing completion of the phage cycle. This is consistent with our observation that four systems (Ogmios, Sucellos, Esos and Belisama) present domains common or homologous in systems causing growth arrest or cell death upon infection (Figure 2).

**FIGURE 3:**
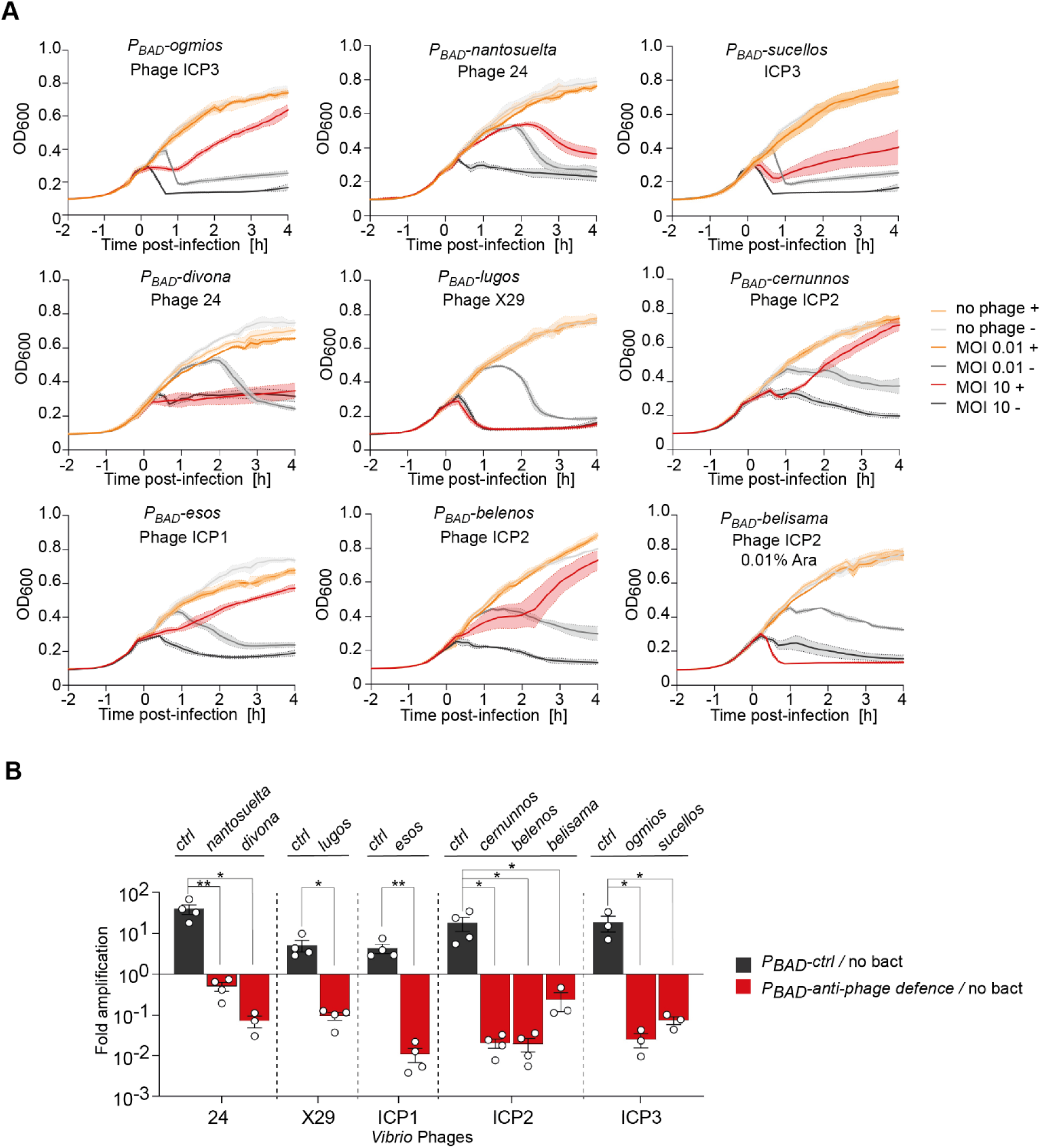
Dynamics of phage infection in presence of anti-phage cassettes. **A.** Growth curves in liquid culture for *V. cholerae* N16961 bacteria that contain the anti-phage system (+) or not (-), infected by phages (MOI 0.01 or 10) or not (no phage). Bacteria were infected at time 0, and measurement of the optical density (OD 600nm) was taken every 10 min. For each experiment, three biological replicates are presented as individual curves. **B.** Amplification of phages in *V. cholerae* strains harbouring a plasmid expressing the anti-phage defences or not (ctrl) or without bacteria (no bact). Fold amplification measured as the ratio between *V. cholerae* strains and no bacteria are shown. Bar charts show the mean fold amplification of at least three independant experiments (n≥3, individual plots) and error bars show the standard error of mean (SEM). Statistical comparisons (Student’s t-test, one-tailed) are as follow: **Pvalue < 0.01 and *Pvalue < 0.5.

We analyzed in detail the Sucellos system with Alphafold (*53*). Our prediction indicates that the SclA and SclB proteins each make a homomultimer with the SclB protein forming a transmembrane pore (Figure S6A). Transmembrane proteins are known to mediate cell suicide in other defence systems (*33, 54*). This cassette contains its own promoter that we named P_sucellos_. To get closer to the natural conditions of expression, we cloned and expressed this system under the control of P_sucellos_. While Sucellos induces a strong resistance against ICP3, we observed smaller plaques using phages 24 and ICP1 but no significant reduction of the number of plaques. This was not previously seen using the inducible P_BAD_ promoter (Figure S6B). Infection in liquid culture and phage amplification assays confirm the suicide mechanism triggered by this cassette this time induced by the phage 24 (Figure S6C). We also performed real time microscopy to visualize cells during phage 24 infection (Figure S6D). Cells devoid of the anti-phage cassette were not protected against the phage and exhibited phage-mediated cell lysis around 30 min after initial infection. However, several cells containing the Sucellos system exhibited lysis right after infection (at 0 min), indicating that in these conditions the Sucellos system induces cell death in infected cells, preventing the completion of the phage cycle and the phage mediated lysis.

To gain insight into the function of the Sucellos system, we performed a phage bacteria co-evolution experiment, and we isolated escapers of the phage 24 able to overcome this system. Whole genome sequencing of 6 escaper phages revealed that all of them carried common mutations (Y112R and N110N/N115D) in the gene *24_60* (Figure S7A), which encodes a pentapeptide repeats containing-protein (Figure S7B). We hypothesized that 24_60 protein might counter the anti-phage activity of Sucellos. We deleted the *24_60* gene of the phage 24 and infected *V. cholerae* with this edited phage. In the presence of the Sucellos system, we obtained a very high fold resistance compared to the infection with the *wt* phage (Figure S7C). We can therefore postulate that the 24_60 protein works as an anti-defence system and that obtained mutations increased its effect against Sucellos.

### Anti-phage cassettes can efficiently recombine and be expressed in the *V. cholerae* SCI

As the majority of integron cassettes are devoid of promotor, their distance from the P_C_ promoter is important for their expression (*55–57*). All the anti-phage cassettes discovered here were found to be located far from the P_C_ promoter in the *V. cholerae* SCI and probably not sufficiently expressed to confer phage resistance. We decided to test the effect of translocating one system, Divona, to the first position of the SCI array. We first deleted this cassette (SCI with Divona deleted, Figure 4A). We then inserted it at the first position in the *attI* site, between the G and T nucleotides of the GTTA cleavage site, exactly where recombination is expected to occur (SCI with Divona relocated). This shuffling event should ensure cassette expression, thanks to the presence of the P_C_ promoter. We then confirmed that integration of the Divona-containing cassette in the first position activated the anti-phage function against phage 24 compared with *wt* or the deleted Divona strains (Figure 4A). As there are a few weak promoters in the integron array (such as those of the TA cassettes, (*23, 58*)), we mimicked an excision event of cassettes located just upstream of the Divona-containing cassette (SCI with Sucellos to VCA0373 deleted) and demonstrated a slight activation of the Divona anti-phage defence, as we observed a decrease in plaque size (Figure 4A).

**FIGURE 4:**
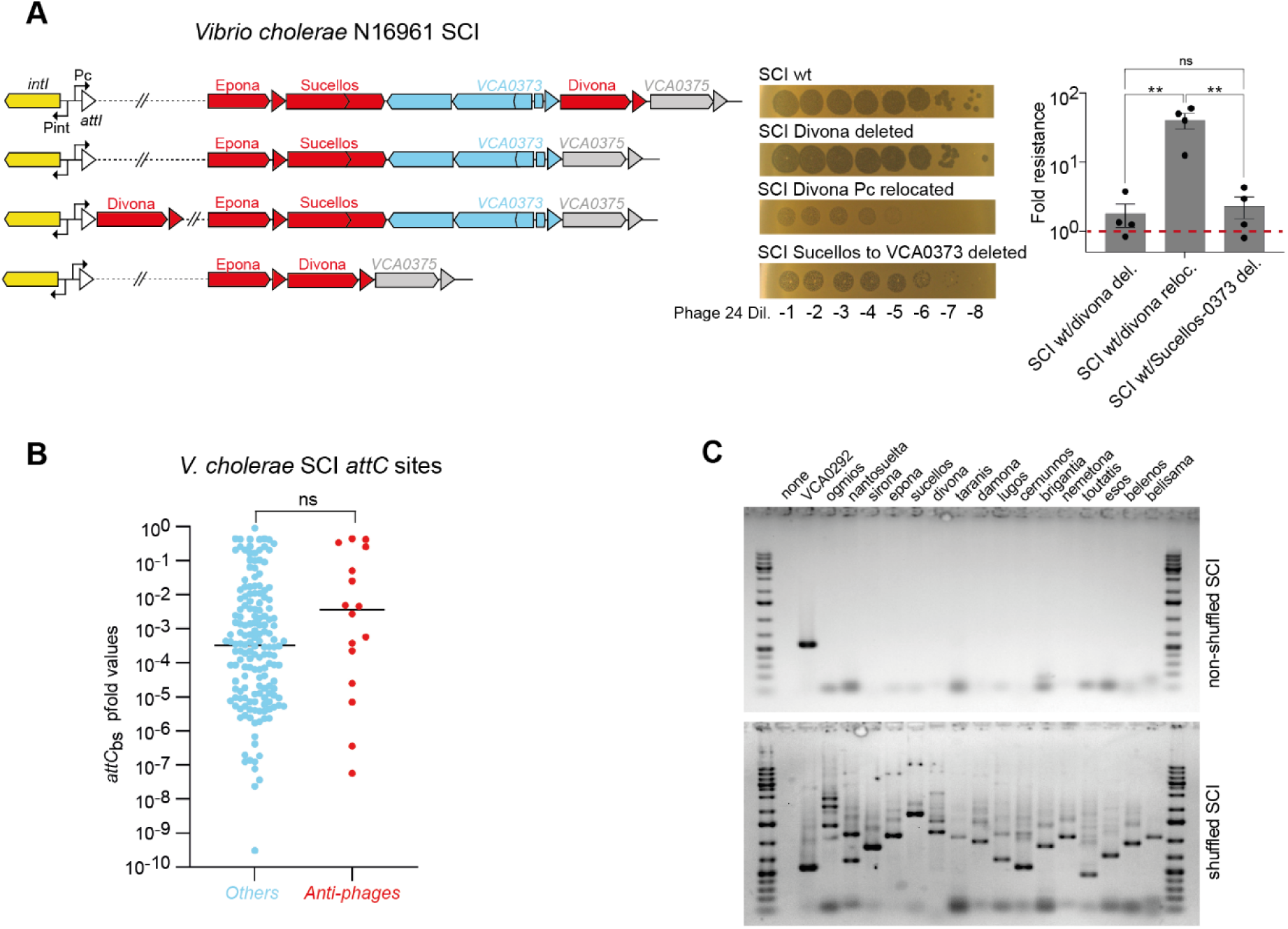
Efficient recombination and expression of anti-phage cassettes in the *V. cholerae* SCI. A. Schematic representation of the *V. cholerae* constructed strains is shown on the left panel. The 10-fold serial dilutions of a high titer lysate of phage 24 spotted on these constructed strains are shown in the middle panel. Representative images of a single replicate out of four independent replicates are shown. Fold resistances measured as the ratio between WT and constructed strain values are shown on the right panel. Bar charts show the mean fold resistance of four independent experiments (n=4, individual plots) and error bars show the standard error of mean (SEM). Statistical comparisons (Student’s *t*-test, two-tailed) are as follow: ns, not significant; \*\**Pvalue < 0.01*. B. Box plot showing the distribution of *attC*_bs_ pfold values of the 179 *attC* sites (bs: bottom strand) of the *V. cholerae* SCI. In red, the 16 *attC* sites associated with the 16 anti-phage cassettes and in blue the remaining (others). Statistical comparison (Mann-Whitney test) is as follow: ns, not significant, *P*=0.1308. C. Results of the PCR made on the library of shuffled cassettes to identify the anti-phage cassette integration in the first position in the array. The non-shuffled SCI strain is used as control. PCRs were performed using a forward primer specific for the beginning of the integron (non-variable part) and a reverse primer specific for each cassette tested. The word “none” means that only the primer specific for the non-variable part is used, and that no cassette-specific primers are used. The multiple bands, seen just above the band of interest only in the shuffled SCI, correspond to the specific shuffled cassette but with one or more other cassettes inserted before it.

The frequency of cassette insertion depends on the properties of *attC* sites involved in the reaction (*59, 60*), which must adopt a folded single-stranded recombinogenic structure to bind the integrase (*59, 61*). We calculated the folding probability (pfold (*62*)) of the 179 *attC* sites of the *V. cholerae* SCI. We found no significant differences between the pfold distribution of the 16 anti-phage cassettes and the others (Figure 4B). This analysis suggests that anti-phage cassettes can be mobilized like the other cassettes. This was confirmed experimentally using a previously constructed inverted SCI strain, which achieves a detectable recombination rate along the entire integron (*17*). We constructed a library of shuffled cassettes by inducing the integrase and used PCR primer pairs specific to each anti-phage cassettes to identify them. We were able to locate all anti-phage cassettes at the first position, even those with a low pfold (that is, Damona-containing cassette, Figure 4B and 4C) meaning that the *attC* sites of anti-phage cassettes are functional. We conclude that *V. cholerae* anti-phage cassettes can recombine and be expressed when positioned in first position in the integron array.

### Frequency and locations of the newly discovered anti-phage defence systems

We searched for homologs of the 16 anti-phage systems across Bacteria to understand their frequency and location. Since most systems have one single gene one cannot use co-localisation between genes to separate true from false positives. Hence, we used blastp (except for *tutA,* searched with tblastn, see Methods) instead of HMM models or protein fold predictions to identify homologs, which may have rendered our search conservative. Among the 2505 identified systems (Table S7), around a third (807) were found in *V. cholerae*. These are usually more than 90% identical to the ones found in *V. cholerae* El Tor str. N16961, suggesting one single ancestral acquisition in the species. A noticeable exception concerns Belisama that is in two copies in two strains, one very similar to the reference and the other very divergent (56% identity in protein) suggesting multiple past events of horizontal gene transfer. This is also the case for other systems found in environmental *V. cholerae* strains. Strikingly all the homologs in the species are found in the SCI, suggesting that they have been part of integron cassettes ever since they were acquired by the species. The median genome has 10 homologous systems, but the detailed analyses of the genomes shows that most have either almost all systems or none. To understand this pattern, we represented the systems in the context of a core-genome phylogenetic tree of *V. cholerae* (Figure 5A). While the systems are scattered across the integron, their relative order is highly conserved, with some arrays presenting cassette deletions and a few presenting translocations. These few events complicate the identification of the exact order of acquisition of cassettes. Yet, it is noticeable from the phylogenetic analysis that most systems were acquired recently at the onset of the emergence of the 6^th^ and 7^th^ pandemic lineages (Figure 5A). This result contrasts with the analysis of previously known defence systems which were more frequent in environmental strains of *V. cholerae* (Figure S1). This suggests a change in the ecology of phage-vibrio interactions when the pandemic strains emerged.

**Figure 5.**
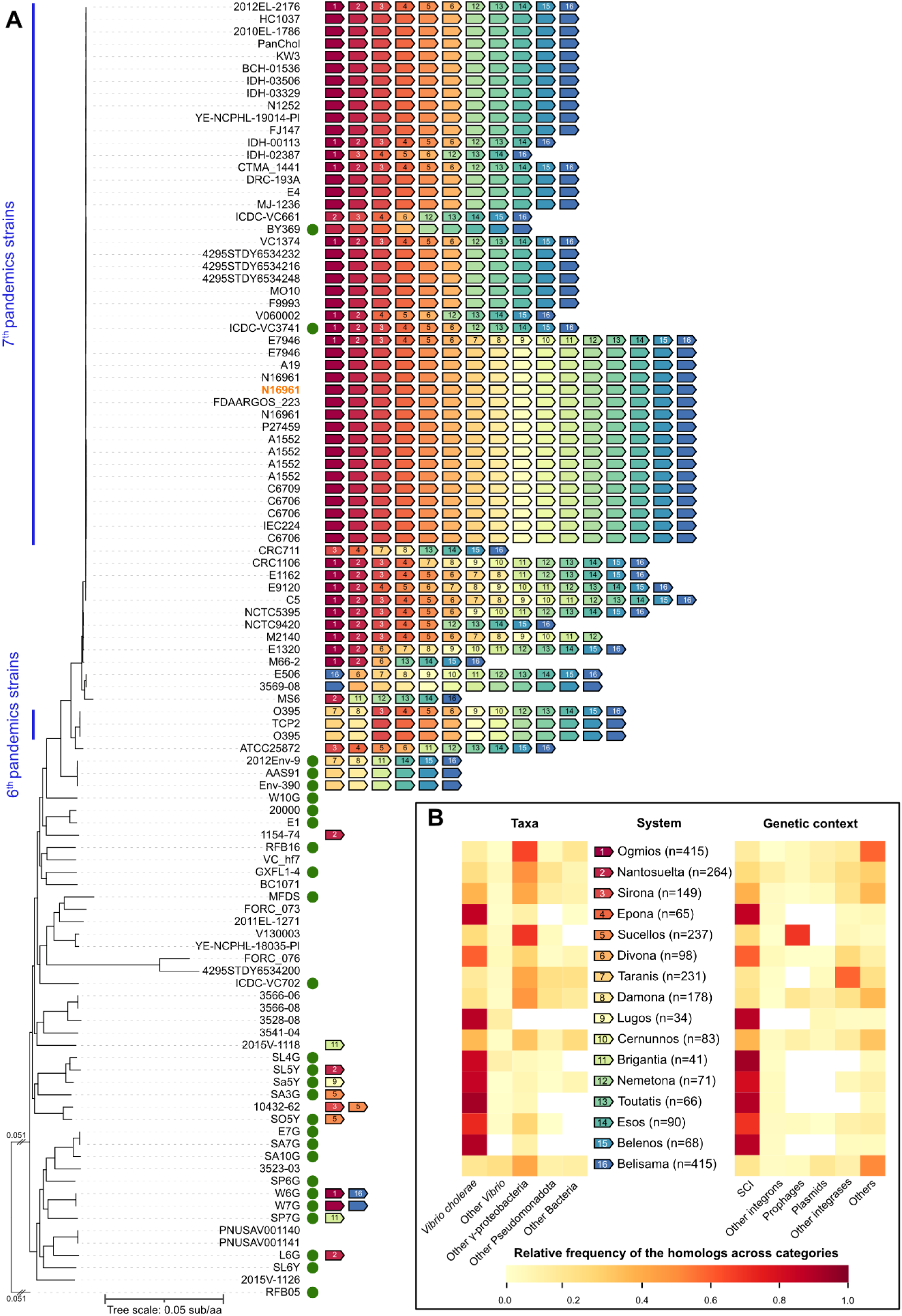
Analysis of the homologs of the discovered anti-phage defence systems. **A.** Presence of the homologs of defence systems in *V. cholerae* plotted in the core genome tree. Defence cassettes are indicated by an arrow and ordered by their position in the SCI. The colour and number give the identity of the cassette and the relative position in the integron of the focal strain *V. cholerae* El Tor str N16961 (indicated in orange in the list of strains, see panel B). The strains from the 6^th^ and 7^th^ pandemic are indicated by a blue vertical line on the left. Green dot: environmental *V. cholerae* Heatmaps of the relative frequency of the discovered defence systems across taxa (left) and genetic contexts (right). The scale of colours is given in the bottom of the graph. The cassettes are indicated as in panel **A**: within the arrow is indicated the relative order of the cassette in the focal strain *V. cholerae* El Tor str N16961 and within parenthesis is indicated the number of homologs.

Several systems are almost only present in *V. cholerae* (Figure 5B), usually not the most abundant ones. Interestingly, some of these showed anti-phage activity only in *E. coli* (e.g., Epona, Toutatis) and, for a given system, we could not find a clear correlation between the proportion of homologs found outside of *V. cholerae* and the specificity of the anti-phage patterns with respect to the two species (as represented in Figure 2A), *i.e.*, the systems restricted to *Vibrio* spp were not necessarily those giving null results in the *E. coli* genetic background.

We then analyzed the taxonomic distribution and genetic context of the two thirds (1698) of homologous systems that were found in other species. While our dataset had 408 genomes of other *Vibrio* spp, the systems were not very abundant in the genus (151 found). Instead, they could be found in the other Pseudomonadota (regrouped for clarity in Figure 5B). The most extreme cases were Belisama, found in 70 genera of Pseudomonadota, Nantosuelta (40 genera) and Sirona (34 genera). Our analyses already confirm that these discovered systems can be found in many other bacterial clades, even in those that were previously found without any integrons (*19*). For example, Cernunnos was found in *Bacteroidota* and Taranis (resembling AbiJ, initially discovered in *Lactococcus lactis*) in Bacillota and Actinomycetota.

We then investigated the genetic context of systems found outside *V. cholerae*. The ones present in integrons were usually found in other species of the *Vibrio* genus (Figure 5B). Outside integrons, the systems were identified in a variety of contexts that differed between systems. For example, several were preferentially identified in prophages, such as Sucellos (the RexAB-like system, often in *Salmonella*), Damona (*Acinetobacter*) and Cernunnos (*Escherichia*). While the unknown anti-phage systems we describe here are rarely encoded on plasmids, some of them are specifically found in chromosomes less than 20kb away from integrases, suggesting they are part of mobile genetic elements (e.g. Taranis, in *Burkholderia* and *Klebsiella;* Divona in *E. coli*) (Figure S8). Hence, these discovered defence systems occur in a variety of genetic contexts. In *Vibrio cholerae*, they are always part of the SCI, whereas in the other species they are often found in mobile genetic elements.

## DISCUSSION

We started this study to test the hypothesis that integrons are biobanks of defence systems. We confirmed that many known defence systems are in integrons. These systems are not a random sample of the types found in genomes. Instead, they tend to be among the smallest. It was previously observed that defence systems in phage satellites are smaller than those on phages, which are themselves smaller than those on other locations of the bacterial chromosome (*63*). This was interpreted as the result of constraints on genome size caused by phage packaging. Here, this trend is pushed to the extreme. This is probably a constraint imposed by the action of the integron integrase, because long cassettes recombine poorly (*11*). The low rates of mobilization of long cassettes defeat the purpose of long loci encoding defence systems being in the integron biobank. Our results reveal an additional crucial characteristics of defence systems harbored by integrons: among homologous groups of defence systems, those in integrons tend to be smaller than those in other locations of the genome. Hence, integron defence systems are small by two synergistic effects: the types of systems (those encoded in one or two genes) and the size of the genes of the defence systems (that are smaller). This suggests the existence of a process of streamlining of molecular systems to fit integron cassettes. As a result, there may be plenty of minimal versions of defence systems in integrons and these could be an untapped source of tools for biotechnology.

We were puzzled by the lack of known defence systems in the integron of our *V. cholerae* strain N16961. We therefore decided to study the large number of cassettes in the integron that were of unknown function. Since genes in integrons are often poorly or not expressed, we cloned each cassette onto a plasmid and found that ca. 18% of the tested cassettes have measurable activity against a panel of phages in *V. cholerae* or *E. coli*, meaning that at least 10% of all *Vibrio cholerae* SCI cassettes encode anti-phage systems. This is probably an underestimate. Some systems may have been missed because they provide defence against other types of phages, might require distinct growth conditions, or may protect from other types of MGEs, e.g. plasmids (*25, 64*). If we take all integrons in our genome database and cluster the proteins encoded in their cassettes at low sequence similarity (30% identity) we obtain almost 25 thousand distinct families of cassettes. If 10% of them are anti-phage systems, integrons may be an extraordinary repository of anti-phage functions.

We have increased the ability to uncover defence systems by using two different genetic backgrounds and panels of phages. Most previous studies cloned defence systems in a model organism (*E. coli*, *B. subtilis*, *Pseudomonas*) and tested them against a panel of phages. We could not find a large-scale study where several systems were systematically tested both in their native host and in a model organism. Here, using the same set of systems expressed from the same plasmid backbone, we observed defence activities for different subsets of systems in the native *Vibrio* strain (9 anti-phages systems revealed using 5 phages) and in *E. coli* (9 anti-phage systems revealed using 23 phages), with little overlap between species. These intriguing differences may be caused by at least three factors. First, the matrices of defences are sparse, i.e. most systems provide resistance to a few phages. This is common in other phage-bacteria models (*41, 63, 65*). Using a larger number of phages might better evidence the systems that have a broader range of activity. Second, defence system expression or activity might be affected by the host genetic background. If so, unknown systems may remain to be identified in the SCI by using more diverse panels of phages and bacterial hosts. Finally, phages and *V. cholerae* strains have a long history of antagonistic co-evolution (*66*), and one cannot exclude that the *Vibriophages* tested in this study already developed anti-defences against some of the integron cassettes. Indeed, we have found one probable anti-defense phage protein protecting against Sucellos (Figure S7).

How do integrons favor bacterial adaptation to phage pressure? We believe this occurs in three ways. First, the integration of a novel phage-resistance cassette in a SCI facilitates the acquisition, stabilization, and expression of a potentially advantageous function under phage pressure. Second, antagonistic co-evolution between phage and bacteria, with phages eventually becoming resistant, is expected to lower the benefits associated with the phage resistance cassette. The natural dynamics of integration of novel cassettes in the SCI increasingly separates the defence system from the integron promotor, thus becoming part of a biobank of non-expressed defence systems. Third, events of recruitment of existing integron cassettes to the expressed position seem more likely (or quicker) than the acquisition of defence system by horizontal gene transfer. If the intrinsic rate of cassette shuffling is high enough, or is stimulated by the presence of a phage, then a bacterial variant where the cassette becomes under the action of a promoter may protect the bacterium from the phage and sweep through the population. Yet, our results suggest such a scenario may not be very frequent in *Vibrio cholerae* because the shuffling rates obtained in laboratory conditions were low (*17*), and we found few rearrangements in the defence cassettes of natural isolates (Figure 5A). Also, systems causing cell death upon infection will not provide effective protection if they are rare in the population. We propose a second scenario that is compatible with the observed abundance of these systems in the integrons and the low rates of integron shuffling. Environmental *Vibrio cholerae* is often associated with host organisms or particulate matter where it may form metapopulations. These sub-populations may occasionally endure cassette shuffling placing a defence system near the promoter of the integron (Figure 6). If the expressed system allows infected cell survival in a diverse population, it may be immediately selected for (Figure 6B). If the system induces death of infected cells (Figure 6C and D), it will provide protection to the population (*33*) only if a fraction of cells containing this system remain uninfected in a homogenous population (Figure 6D). The fixation of cell death inducing mechanisms is therefore more difficult and may require population structure as provided by biofilms, processes of host infection, or colonization of particles. To test these hypotheses, we will need a better understanding of the dynamics of gene cassettes, their impact on the expression of the defence systems, and how they affect selection for anti-phage functions in the integron.

**FIGURE 6:**
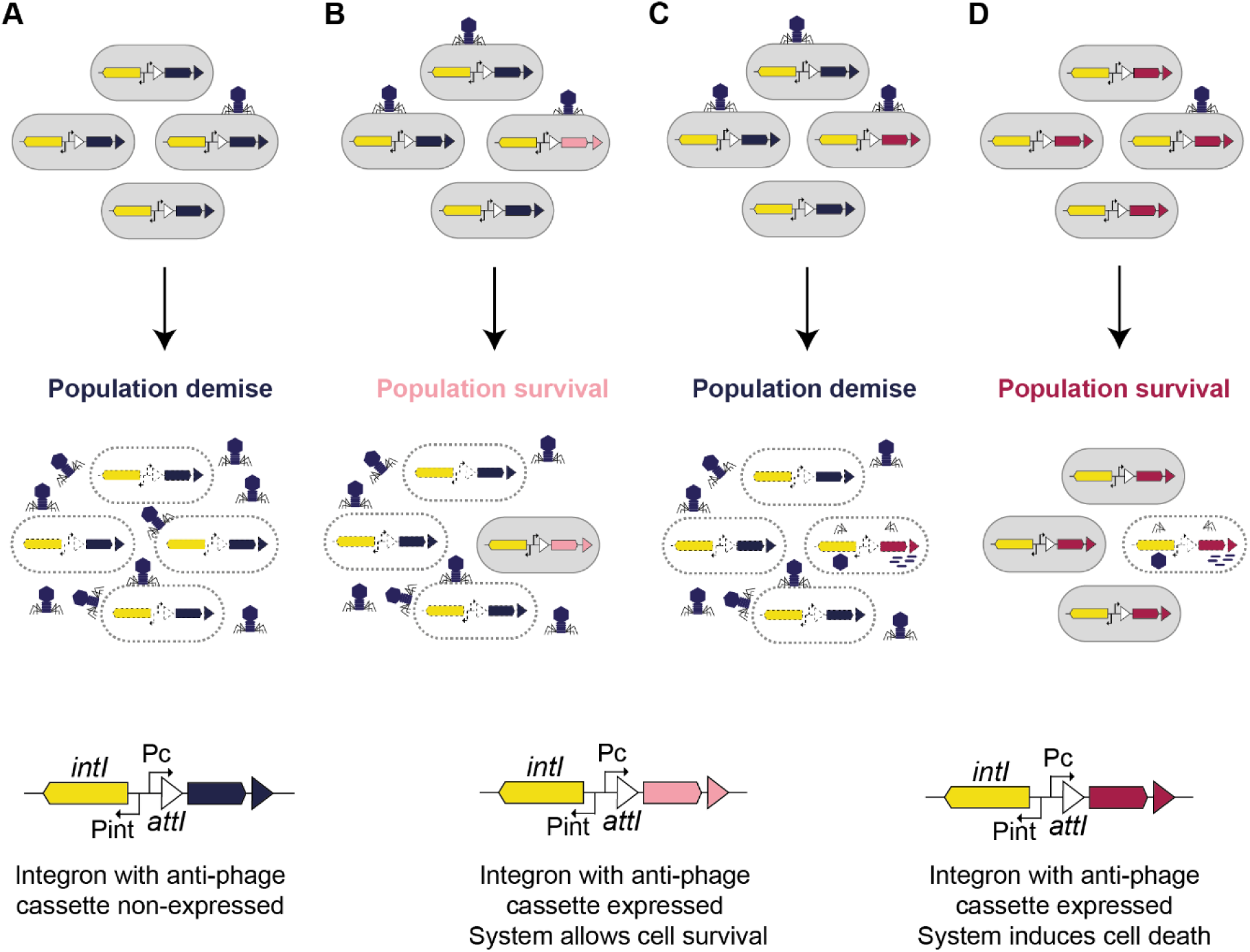
Integron cassette shuffling and bacterial adaptation to phage pressure. **A.** Any cell in the population expresses anti-phage cassette, the population is lysed and the phage is amplified. **B.** Only a small fraction of the population expresses an anti-phage cassette that allows cell survival, the population survives. **C.** Only a small fraction of the population expresses an anti-phage cassette that induces cell death, the population is lysed and the phage is amplified. **D.** The whole population expresses an anti-phage cassette that induces cell death, the population survives at condition that only a part of the population was infected.

Past work on the function of integron cassettes has focused on antibiotic resistance, which is frequent in mobile integrons, but rare in SCIs. Hence, the function of the vast arrays of SCI cassettes has remained an enigma for decades. Here, we unveil a large set of functions encoded by these elements. As cassettes in SCIs can be mobilized to MIs and then spread by horizontal gene transfer, they may drive resistance to phages across Bacteria as shown by a simultaneous study by Kieffer *et al* (*bioRxiv*). It is possible that many of the other functions of these integrons are also unknown defence systems. The identification of SCI as a hotspot of anti-phage systems has similarities and differences with the defence islands whose study led to the discovery of so many unknown defence systems (*46, 67*). Both regions concentrate anti-phage systems, but defence islands seem to have originated by successive integration and excision of mobile genetic elements, suggesting that at least initially the systems were selected for the maintenance of the MGE, which may or may not favor the host (*68*). In contrast, SCIs are not part of MGEs and are not mobile *per se*. Their anti-phage systems are the result of the creation or exchange of cassettes between integrons, and their subsequent stabilization strongly suggests they are specifically adaptive to the host.

Integron with any anti-phage cassette expressed is represented at bottom left of the figure. The expressed non anti-phage cassette is represented in blue. The same integrons but with anti-phage cassettes expressed (i.e. recombined in first position of the array), are represented at the bottom right of the figure. Expressed anti-phage cassettes that allow cells to survive are shown in light pink and those that induce cell death in dark pink.

## METHODS

### Bioinformatics analysis

#### Data

Except when specifically mentioned, the genomic analyses were done with a dataset of 32,798 completely assembled genomes from the NCBI RefSeq database of reference microbial genomes (downloaded on May the 15^th^ of 2023). The full list is available in Table S1.

#### Identification of integrons in complete genomes

We used the integron predictions obtained by IntegronFinder 2.0.5 (*19*) on the RefSeq database (default options, –local-max for more sensitive detection and –gembase to preserve annotations). The complete list of these predictions can be found in Table S3. For each genome, integrons were detected replicon by replicon. IntegronFinder provides three subtypes of integron-like elements: complete integrons (IntI plus a few cassettes flanked by *attC* sites), CALINs (Cluster of *attC* sites lacking an integrase) and loner integrases. In the current study, we considered every cassette protein belonging to integrons (67% of total) or CALIN detected by IntegronFinder.

#### Identification of defence systems

We searched for defence systems in all of the genomes of the database using DefenseFinder v1.2.0 (*69*), with the –db-type gembase and –preserve-raw options, and the DefenseFinder models database v1.2.4 (downloaded on February the 14^th^ of 2024). This model database comprises custom HMMs (Hidden Markov Models) of the components of 152 experimentally validated defence systems that are searched for at the same time. A defence system is identified when all its mandatory components are found co-localized in the genome. The option –db-type gembase allows to test for this co-localization at the replicon level.

#### Integration of the results of IntegronFinder and DefenseFinder

The predictions of IntegronFinder and DefenseFinder were merged and analyzed jointly to pinpoint defence systems encoded in integrons. Our analysis deems cassettes as autonomous components of the integron, i.e. it assumes that systems are encoded on one single cassette. Indeed, most systems (783/789) were identified in one single integron cassette, i.e. the region between two consecutive *attC* sites. Six systems were found to be split between neighboring integron cassettes. This is unexpected, since cassette shuffling is likely to disrupt this genetic organization and disrupt the system. Hence, we decided to discard these six systems. Similarly, we did not search systematically for components encoded in distant cassettes that could eventually produce a complete system. Likewise, we did not look for systems that could be split between an integron and another part of the genome.

#### In-depth analysis of defence systems in Vibrio cholerae

We made a detailed analysis of the 107 complete genomes of *Vibrio cholerae* in the database above. Strains were classified as either environmental or clinical based on the information available on RefSeq and in the literature. A pangenome, a core genome, and a phylogenetic tree were built using the serial modules of PanACoTA v1.4.1 (*70*). Briefly, we computed the species’ pangenome using mmseqs2 v15-6f452 (*71*) and using a threshold of at least 80% amino acid identity and 80% coverage among genes belonging to the same pangenome family. Then, the species’ persistent genome was defined as the set of pangenome families having a unique member in at least 90% of the genomes. We excluded two strains (*Vibrio cholerae* 2740-80, GCF_001683415.1; *Vibrio cholerae* strain PS4, GCF_019504425.1) that harbored less than 90% of the species’ persistent genome and were hence too distant to the other strains in the species. Using MAFFT (*72*), we aligned individually each pangenome family belonging to the persistent genome (alignments in proteins backtranslated to codons). Finally, the resulting codon alignments were concatenated, and the phylogenetic tree was inferred with IQTree v 2.2.2.2, using the options ModelFinder and computing 1000 ultrafast bootstrap (*73*). The resulting tree was visualized with iTOL (*74*).

### Identification of homologs of the discovered defence systems

We searched for homologs of the discovered defence systems in the RefSeq database using blastp (minimal coverage of the query sequence 50% and eval< 10^-5^). We only made an exception for *tutA* because we realized that this small gene was often not identified in the annotation files. In this case, we used the protein of the focal strain and used tblastn to identify homologs directly in the genome nucleotide sequence (same criteria). The presence of a homologous system when the focal one had 2 genes required the identification of co-localized homologs for both genes. The dGTPase was too frequent and difficult to distinguish between homologs with different functions (*34*). Hence, we sampled among the hits those with eval<1e-73 and coverage of the query sequence 70%. These thresholds allow to select for homologs that do not hit the current DefenseFinder model for dGTPases (Figure S8). A similar problem was identified with the MTase, all of them hitting the DefenseFinder HMM for RM MTases, but in this case the requirement for co-localization helped categorizing the hits.

We aimed at describing the genetic context in which the homologs were found. To detect integrons and related elements, we relied on the IntegronFinder v2.0.5 predictions (same parameters as above, except for --calin-threshold 1 to also detect SALINs (*75*)). We distinguished SCIs (complete integron or CALIN with >10 cassettes) from other types of elements detected by IntegronFinder, grouped into “Other integrons”. Prophages were detected with geNomad v1.5.2 (*76*). Replicons were classified as plasmids based on the RefSeq metadata. Integrases were identified using the PFAM profiles PF00589 for tyrosine recombinases, and the pair of profiles PF00239 and PF07508 for serine recombinases (*77*). All the protein profiles were searched using hmmsearch from the HMMer suite v.3.3.2 (<u>hmmer.org</u>, default parameters). Hits were regarded as significant when their e-value wassmaller than 10−3 and their alignment covered at least 50% of the protein profile. The complete list of these homologs can be found in Table S7.

### System description

We investigated whether the newly identified systems had protein domains found in known systems using available data (Uniprot, (*29*)), sensitive sequence similarity detection programs (HHPred (*30*)), and structural similarity (AlphaFold (*31*) and FoldSeek (*32*)). All of the relevant information is available in Figure S4.

### AlphaFold structure prediction

Alphafold-Multimer (*53*) was used to model the structure of complexes. The predictions were run via ColabFold v1.5.5 (*78*) using default parameters.

### Bacterial strains, plasmids and primers

The different virus and bacteria strains, plasmids and primers that were used in this study are described in Table S8, S9, S10 and S11.

### Media

*Escherichia coli* and *Vibrio cholerae* were grown in Luria Bertani lennox (LB) at 37°C. In the case of *E. coli*, antibiotics were used at the following concentrations: carbenicillin (Carb), 100 µg/ml, kanamycin (Km), 25 µg/ml, spectinomycin (Sp), 50 µg/ml. Diaminopimelic acid (DAP) was supplemented when necessary to a final concentration of 0.3 mM. To induce the P_bad_ promoter, L-arabinose (Ara) was added to a final concentration of 2mg/ml; to repress it, glucose (Glc) was added to a final concentration of 10mg/ml. To induce the P_tet_ promoter, anhydrotetracycline (aTc) was added to a final concentration of 100 ng/ml. *V. cholerae* strains were cultivated in the same conditions and with the same antibiotic concentrations except for Sp, that were supplemented at a final concentration of 100 μg/ml. When *V. cholerae* strains were cultivated in presence of glucose, the later concentration of Sp was increased 2-fold (200 µg/ml). CaCl_2_ and MgSO_4_ were used at a final concentration of 10 mM for *V. cholerae*, and CaCl_2_ at a final concentration of 5 mM for *E. coli*.

### Strain and plasmid constructions

All PCR reactions were performed using the Phusion High-Fidelity PCR Master Mix (Thermo Scientific, Lafayette, CO, USA), and all diagnostic PCR reactions were performed using DreamTaq DNA Polymerase (Thermo Scientific, Lafayette, CO, USA). Oligonucleotides were synthesized by Eurofins Genomics. Gibson Assembly was performed using the master mix (New England Biolabs, NEB) according to the manufacturer’s instructions. Genomic DNA was extracted using DNeasy® Tissue Kit (Qiagen) and Plasmid DNA using GeneJET Plasmid Miniprep kit (Thermo Scientific, Lafayette, CO, USA). Natural transformation was performed according to the procedure described in (*79*).

### Cloning of the SCI cassettes in the pSC101 expressing vector

Among the 179 cassettes of the SCI, we excluded those of known function, those coding toxin-antitoxin genes (TA cassettes), and the second copies of duplicated cassettes bringing us down to 88 cloned cassettes. Each cassette fragment (corresponding to the 88 cassettes and *tutA*, *tutB*, *sclA* and *sclB* CDSs alone) was amplified from 8637 with corresponding .for and .rev primers. The pMP180 vector was amplified by inverse PCR with 6705 and 7086 primers. Assembly of each two fragments was achieved by performing Gibson Assembly.

### Phages

ICP1, ICP2 and ICP3 phages (for the International centre for Diarrhoeal Disease Research, Bangladesh <u>C</u>holera <u>P</u>hage) represent the three *V. cholerae*-specific, lytic (virulent) classically used phages (*80*). They were isolated from rice-water stool samples of cholera patients in Bangladesh (*66*). ICP1 and ICP3 receptors were identified as the lipopolysaccharide (LPS) O1 antigen (*81*) and that of ICP2 as the major outer membrane porin OmpU (*82*). The two other phages (24 and X29) were transmitted from the *Vibrio cholerae* CNR of Institut Pasteur. These phages of our *vibriophage* collection encompass 5 of the 9 *vibriophage* genome clusters identified by Barman et al (*83*).

### Phage amplification

Phages 24 and X29 were amplified using HER1051 as indicator strain and phages ICP1, ICP2 and ICP3 were amplified using MAK757. Single phages from mother stocks were used to prepare working stocks. Mother stocks were streaked into soft agar mixed with culture of the indicator strains and incubated overnight at 37°C. The next day, single plaques were cored with a cone and resuspended overnight at 4°C under agitation in 1mL of LB to let the phages diffuse in the medium. The resulting lysates were then serially diluted and mixed with bacteria at OD_600_ 0.1 grown in LB + CaCl_2_ + MgSO_4_. Once lysed, cultures were centrifuged at 3000 g for 5 minutes and supernatants were filtered through a 0.2 μM membrane. Lysates were then titrated by drop assay and the ones with the highest titers were selected.

Coliphages were amplified on *E. coli* K-12 MG1655. Phage stocks were amplified by mixing 200 μL of an overnight culture of *E. coli* with 10 μL of phage stock solution (either pure or diluted) a or a piece of frozen phage and 25 mL of warm (∼50°C) LB + CaCl2 + 0.5% agar and poured on LB + CaCl2 + 1% agar plates. Plates on which confluent lysis was observed were used to recover the top-agar layer in 1mL of PBS and transfer it to a 50 mL conical tube. The top agar was then disrupted by vortexing until broken into small pieces. Tubes were then left to incubate 10 min at room temperature before centrifugation at 3,000 g for 5 min. Finally, the supernatant was recovered.

### Phage plaque assays

#### Preparation of V. cholerae soft agar plates

Strains carrying each of the systems or the control plasmid were grown overnight in LB + Carb + Glc. The cultures were then diluted 1/100 in LB + Carb + Ara and grown overday up to OD_600_ 0.3. Bacterial lawns were prepared by mixing 100 μL of overday culture with 4 mL of melted LB 0.5% agar + Carb + Ara + CaCl_2_ + MgSO_4_ and the mixture was poured onto round plates of LB 1.5% agar + Carb + Ara + CaCl_2_ + MgSO_4_.

#### Preparation of E. coli soft agar plates

Strains carrying each of the systems or the control plasmid were grown overnight in LB + Carb + Glc. Bacterial lawns were prepared by mixing 200 μL of overnight culture with 25 mL of melted LB 0.7% agar + Carb + Ara + CaCl_2_ and the mixture was poured onto squared plates of LB 1.5% agar + Carb + Ara + CaCl_2_.

Serial dilutions of high-titer (>10^8^ pfu/mL) stocks of phages were spotted on each plate and incubated overnight at 37°C. The next day, plaques were counted, and the fold resistance was measured as the number of plaques in the control plate divided by the number of plaques in the presence of each system. When plaques were too small to be counted individually, we considered the most concentrated dilution where no plaque was visible as having a single plaque.

### Time course infection experiments

*V. cholerae* strains carrying each of the systems or the control plasmid were grown overnight in LB + Carb + Glc. The cultures were then diluted 1/100 in LB + Carb + Ara and grown for two hours. OD_600_ was measured and culture concentration normalized, then diluted again 1/100 in LB + Carb + Ara + CaCl_2_ + MgSO_4_. Note that for Belisama, we used a lower Ara concentration (i.e. 0.01% instead of 0.2%) due to the observed gene toxicity. 180 μL of each preparation was distributed by triplicate in a 96-wells microplate and growth was monitored on a TECAN Infinite M200Pro microplate reader at 37°C with on maximum agitation with absorbance measurements (600 nm) taken at 10-min intervals. When OD_600nm_ reached mid-log phase ∼0.3, cells were either kept uninfected or infected with MOI ∼0.01 or MOI ∼ 10 of phages. Growth was then monitored for 4 h post-infection, with the caveat that the first measurement following infection is always aberrant and was not displayed. Curves correspond to the mean of at least three technical replicates and the shade corresponds to the standard errors at each timepoint. Lysates were also collected 90 min after infection of liquid cultures at initial MOI of 10, and phage titer was quantified by plating serial dilution of the lysates on the strain containing the control plasmid as described above (i.e. Phage plaque assays).

### Microscopy setup for live imaging

*V. cholerae* overnight cultures of the indicated strains were diluted 1/1,000 and then grown in liquid LB + Sp. When OD_600nm_ reached mid-log phase ∼0.3, cells were either infected or not with MOI ∼ 5 of phages and immediately immobilized on 1.4% agarose-padded slides containing the growth medium. A coverslip was placed on top of the agarose pad and sealed with a vaseline:lanolin:paraffin mix (ratio 1:1:1) to prevent pad evaporation. Slides were incubated at 37°C under an inverted wide-field microscope (Zeiss ApoTome) for time-lapse video recording. Video frames were taken at 5 min interval for a total duration of 60 min, using a Plan Apo 63× objective (NA = 1.4) and a Hamamatsu sCMOS ORCA-Flash 4.0 v3 (Pasteur Institute Imaging Facility Imagopole). A total of 3 movies per condition-were recorded and analyzed with FIJI software (*84*).

### *attC* sites pfold predictions

The calculation of the probability of an *attC* site to fold into a recombinogenic structure (pfold) were performed as described in Vit *et al* (62) (*i.e.,* using the RNAfold program from ViennaRNA Package (*85*) (http://rna.tbi.univie.ac.at, category “RNAfold Server”).

### Library cassette shuffling assay

A thermosensitive plasmid expressing the integrase under the control of a TetR-pTet promoter (pR669) was transformed in three SCI Inv *ΔIntIA V. cholerae* replicates. For each replicate, four clones were picked and grown overnight in LB + Sp at 30°C, then diluted 1/1000 in LB + Sp + aTc and grown overnight at 30°C to express the integrase and shuffle the cassette array. Shuffled cultures were diluted 1/100 in LB and grown overnight at 37°C twice to cure the plasmid. Cultures were then streaked on LB agar plates and 16 single clones were picked for each. *attIA* site region was amplified by PCR (using 5778 and 1401 primers) on these clones to detect individual insertion events and validate the shuffled status of our bank. Shuffled cultures displaying cassette insertions in at least 50% of individual clones were then pooled to limit the risk of bias due to an early insertion in of culture. To test the capability of shuffling of the 16 anti-phage cassettes in 1^st^ position in the bacterial library population, we performed PCR using the 5778 primer associated with each cassette specific primer (308.for, 322.for, 356.for, 366.for, 367-368.for, 374.for, 396.for, 399.for, 409.for, 410.for, 419.for, 441.for, 446-447.for, 450.for, 457.for and 458.for). To control the presence of the 1^st^ cassette (VCA0292), we used the 292.for. Each band was sequenced using the 5778 primer to confirm the specific shuffling event.

### Phage training

Phage training was performed according to a method inspired by the Appelmans protocol (*86*).. Phage training was started by mixing early exponential phase *V. cholerae* ΔSCI Δ*ddmABC* expressing or not *sucellos* with 12 different serial dilutions of phage 24 (2 × 10^9^ PFU in the first well). The co-cultures were performed in a 96-well plate, at 180 rpm and at 37°C. After 24h incubation, all wells showing lysis were pooled along with the first non-lysed well past the point of lysis. The pool was centrifugated to eliminate remaining bacteria. The resulting lysate was used for the next round and the process was repeated 5 times (i.e. 6 rounds in total) up to the evolving phage 24 was able to lyse at the smallest concentration. Escaper phenotype was confirmed by plaque assays and phages from 6 different lineages were isolated by picking into single plaques.

### Phage DNA extraction

*Wt* and escaper phages were amplified as described above to obtain high titer stocks (>10^8^ pfu/mL). Aliquots of 500 μL were treated by TURBO DNAse (Thermo Fisher Scientific) and RNAse A (Qiagen) for 1 hour at 37°C to degrade cellular nucleic acids. DNAse was inactivated with the addition of 20 μL of EDTA 0.5 mM. Aliquots were then treated with 0.5 mg/mL of proteinase K (Qiagen) and SDS 0.5% for one hour at 56°C to degrade phage capsids. DNA was then purified as follows: 500 μL ml of a phenol-chloroform-isoamylalcohol (PCI, 25:24:1) solution (Sigma) was added to the sample which was then vortexed and centrifuged for 10 minutes at 16,000 g. The upper aqueous phase (around 400 μL) was transferred to a fresh tube containing 400 μL of chloroform-isoamylalcohol (24:1). The sample was further vortexed and centrifuged for 10 minutes at 16, 000 g, and the upper aqueous phase (around 250 μL) was transferred to a fresh tube containing 25 µl of 3 M sodium acetate. The sample was mixed with 2.5 volumes of cold 100% ethanol and incubated for one hour at -20°C to precipitate DNA. After centrifugation (1 min at 16,000 g), the DNA pellet was washed with 70% cold ethanol. Finally, the pellet was air-dried and resuspended in 50 μL of distilled water.

### Phage sequencing and analysis

Phage DNA was prepared and sequenced at the P2M, Institut Pasteur, using a Nextera XT DNA library preparation kit and the NextSeq 500 System WGS. Trimming and clipping were carried out using *AlienTrimmer* v2.0 with the following parameters: “-q 15 -l 50 -p 50” Trimmed reads of *wt* phage 24 were mapped using bowtie2 (*87*) on the 24 reference genome to verify it. As the repeated region in *24_60* gene was difficult to assemble, it was amplified by PCR with primers 7423 and 7424, and verified by Sanger sequencing. Mutations were only found in *24_60* gene and a new reference genome was generated. This new reference was used to analyze mutations in escaper phages using bowtie2 and IGV browser (*88*). Mutations in *24_60* gene were verified by Sanger sequencing.

### Golden Gate Assembly

The crRNA spacer was introduced into the pBA559 Cas13 expressing vector through a “Golden Gate Assembly”. The spacer was ordered as two complementary primers targeting the transcript of the gene 24_ 60 of the phage 24 and, with 4 bp 5′ overhangs matching the BsaI-digested destination plasmid staggered ends. For the phosphorylation, 6µL (10 µM) of each 7419 and 7420 primer is mixed with 2µL of T4 DNA Ligase Buffer (10X) and 0.4 µL of T4 PNK for a total volume of 20 µL and then incubated for 30 min at 37°C. To insert the guide into the pBA559 vector, we prepare the following mix: 40 ng of the pBA559 plasmid, 2 µL of annealed oligos, 1 µL of Cutsmart buffer, 1 µL of BsaI enzyme, 1 µL of ATP (1mM), 1 µL of ligase, 2 µL H_2_0. We then incubate the mix in a thermocycler with the following steps: 3 min at 37°C (for digestion), 4 min at 16°C (for ligation) and alternate between those steps 25 times. We finish with one cycle of 5 min at 50°C and 5 min at 80°C for enzymes inactivation. At this step, the mix is ready to be transformed in a cloning *E. coli* Top10 strain (Invitrogen) and the successful sgRNA insertions are screened by sequencing using the 7421 primer.

### Two-step phage editing and phage enrichment

To delete the gene *60* of the phage 24, we used the two-step phage editing and phage enrichment method developed by Adler et al, (*89*). First, the phages infect homology vector-containing *V. cholerae* strain at a low MOI, producing a mixed population of original and edited phages. Second, this phage population is diluted and infects a Cas13a-expressing *V. cholerae* strain targeting the original phage locus, enriching the edited phages relative to original. CRISPR-Cas13a proteins are RNA-guided RNA nucleases that bind to complementary target phage transcripts and ensure general, non-specific RNA degradation of infected cells.

#### Phage editing

To create genome-edited phage 24 lysate, the phage-editing strain (W303) containing the homologous recombination vector (pJET-24_60) was grown overnight in LB at 37 °C. The strain was diluted into fresh media (LB + Carb) to an OD_600nm_ of 0.04 or 0.4 in a 96-well plate. Wildtype phage 24 was added to each well to achieve an MOI of 1, 0.1 or 0.01. Infection was monitored in a TECAN shaking at 37 °C at 180 rpm, with OD_600nm_ readings every 5 min. Infection was allowed to proceed until there was a visible population crash (∼7 hours). The well showing a lysis were pooled, centrifuged and filtered on 0.45 μM membrane. These lysates comprise a mixture of homologous recombination-edited phage and wildtype phage.

#### Edited phage enrichment

To enrich genome-edited phage lysates, a phage counterselection strain consisting of W303 containing an ‘enrichment’ Cas13a vector (pBA559-24_60) was grown overnight in LB + Cm media at 37 °C. 100 µl of saturated overnight culture was mixed with molten LB top agar (0.7%) supplemented with Cm and poured on a LB agar plate containing Cm. Dilutions of phage 24 were then spotted and the plates incubated at 37°C. Plaques were isolated and purified for three rounds using the counterselection strain to ensure thorough removal of any remaining wildtype phages. Lysates were centrifugate to remove the remaining bacteria. These lysates comprised an enriched mixture of homologous recombination-edited phage. PCR and sequencing using 7326 and 7327 primers (located outside the homologous region) were performed on phages to confirm the *24_60* gene deletion.

### Statistics and Reproducibility

All experimental data are representative of the results of at least three independent biological repeats. All replication attempts were successful. Statistical analysis of experimental data was done using GraphPad Prism 9.1.2 (GraphPad Software, Inc., CA, USA). Statistical analyses of the computational data were done in JMP (mixed models) and Python (the rest).

## ACKNOWLEDGEMENTS

We thank Florian Tesson for his technical guidance on the analysis of newly discovered anti-phage systems. We gratefully acknowledge François-Xavier Weill and Caroline Rouard from the Centre National de Référence des Vibrions et du Choléra (Institut Pasteur) from the gift of *Vibrio* phages. We gratefully acknowledge the UtechS Photonic BioImaging (Imagopole), C2RT, Institut Pasteur, supported by the French National Research Agency (France BioImaging; ANR-10–INBS–04; Investments for the Future). This work used the computational and storage services (TARS cluster) provided by the IT department at Institut Pasteur, Paris. A CC-BY public copyright license has been applied by the authors to the present document and will be applied to all subsequent versions up to the Author Accepted Manuscript arising from this submission, in accordance with the grant’s open access conditions.

## AUTHOR CONTRIBUTIONS

B.D, M.B, F.D, W. S, K.D, P.A.K and C. L conceived the experimental project, designed the experiments, constructed strains and plasmids, performed the phage assays and analyzed the results. J.B performed Microscopy experiments. E.L and E.P.C.R conceived the computational project, designed the analyses, and analyzed the results. E.L made the bioinformatics analysis with the help of M.T. C.L, B.D, E.P.C.R, E. L wrote the paper with input from the other authors. C.L, D.M and E.P.C.R acquired funding. C.L, D.M, E.P.C.R and F.L.R supervised the project.

---

**Figure S1:**
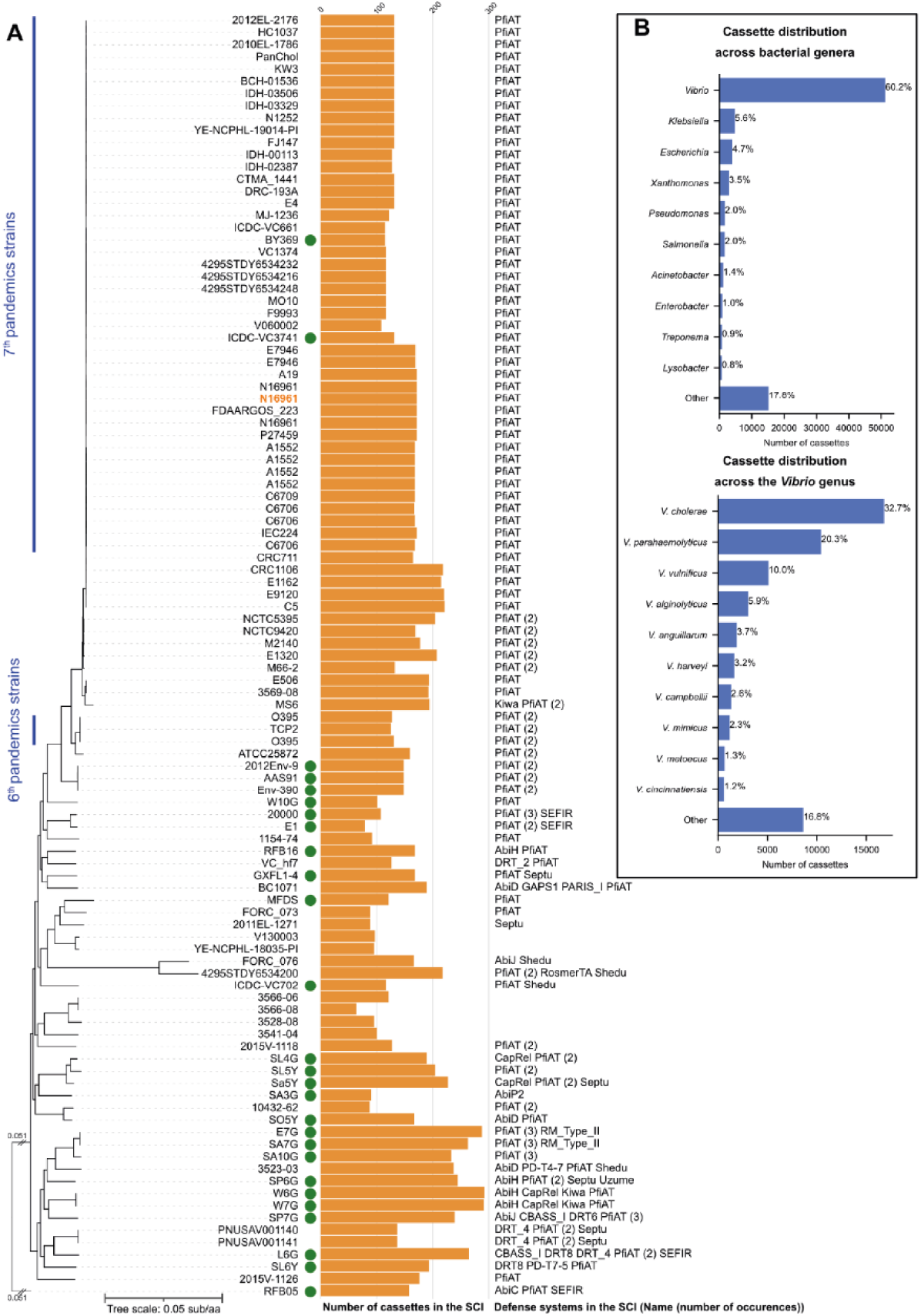
Detailed analysis of the known anti-phage defence systems carried by integrons. **A.** Presence of the known anti-phage defence systems (as detected by DefenseFinder) in the SCI of V *cholerae*, plotted in the core genome tree. The orange barplot shows the number of cassettes (number of affC) carried by each SCI. The names and number of occurences of the systems detected in the SCI are mentioned on the right side. This study’s focal strain *V. cholerae* El Tor str N16961 is indicated in orange in the list of strains. The strains from the 6^th^ and 7*h pandemic are indicated in a blue vertical line on the left. Green dot: environmental *V. cholerae* Distribution (by genus) of the cassettes detected by IntegronFinder in all the complete reference genome database RefSeq (upper panel) / in the *Vibrio* genus (lower panel)

**Figure S2:**
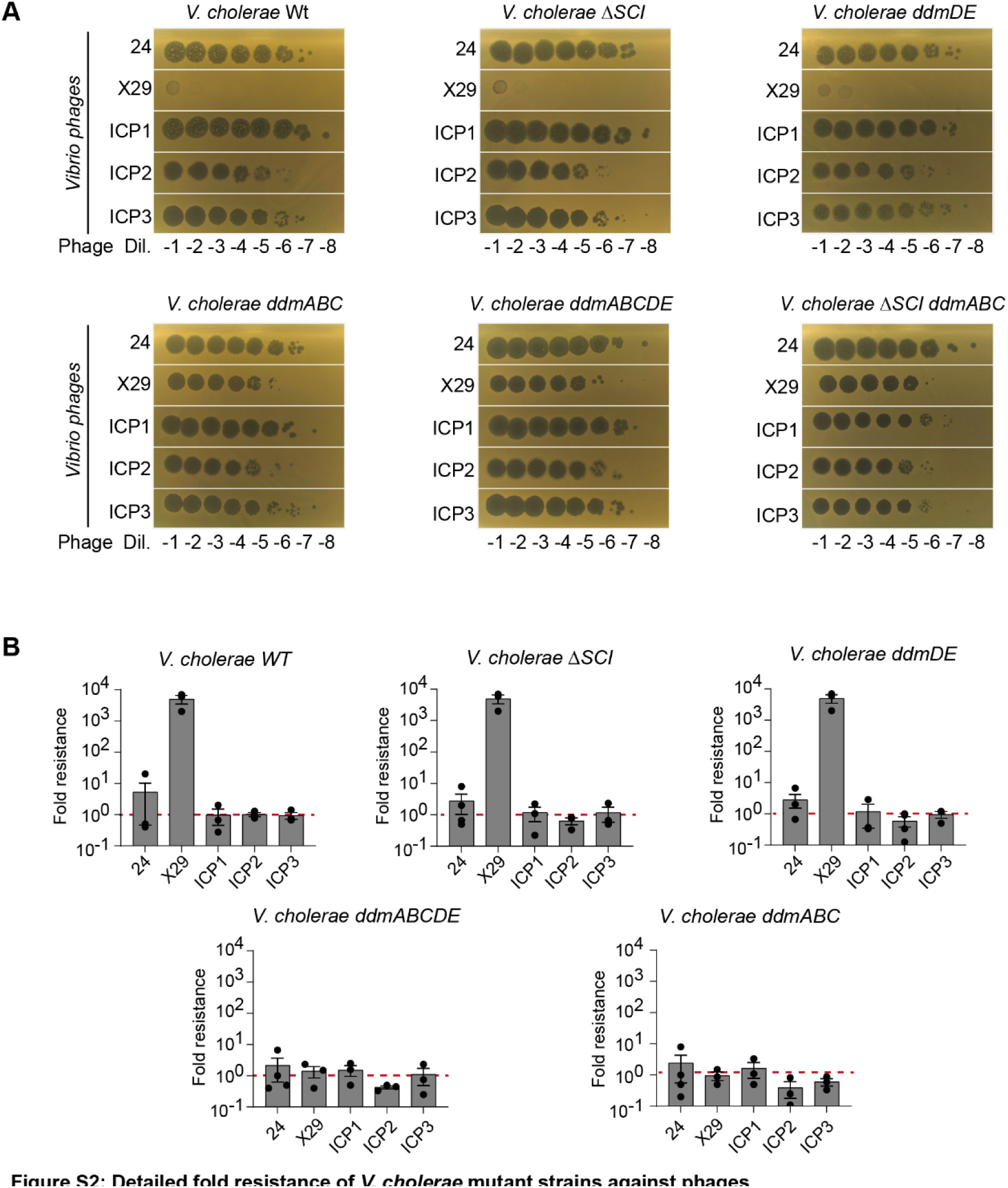
Detailed fold resistance of *V. cholerae* mutant strains against phages. **A.** 10-fold serial dilutions of a high titer lysate of phages spotted on *V. cholerae* strains. Representative images from a single replicate out of three independant replicates are shown. **B.** Plaque-forming units were measured for each phage on the indicated *V. cholerae* strains and on the *ASCI ddmABC* cells. Fold-resistance was measured as the ratio between these two values. The bar charts show the mean fold resistance for three independent replicates (n=3, individual plots). Error bars shown the standard error of mean (SEM).

**Figure S3:**
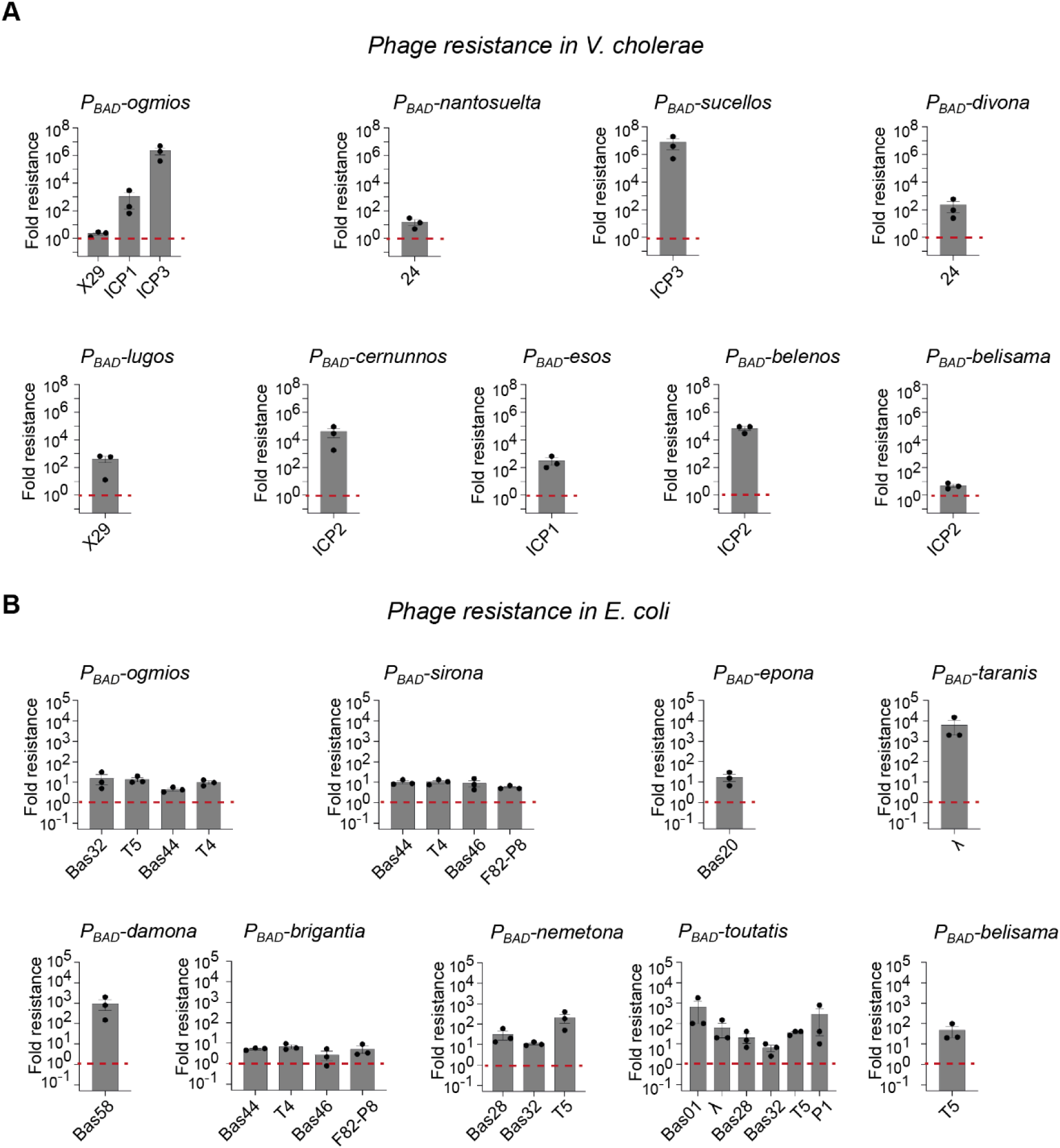
Detailed fold resistance of all defence systems against phages, Related to Figure 2. Plaque-forming units were measured for each phage on *V cholerae* (A) or *E. coli(B)* cells harboring either a control plasmid or a defence system. Fold-resistance was measured as the ratio between these two values. The bar charts show the mean fold resistance for three independent replicates (n=3, individual plots). Error bars shown the standard error of mean (SEM).

**Figure S4:**
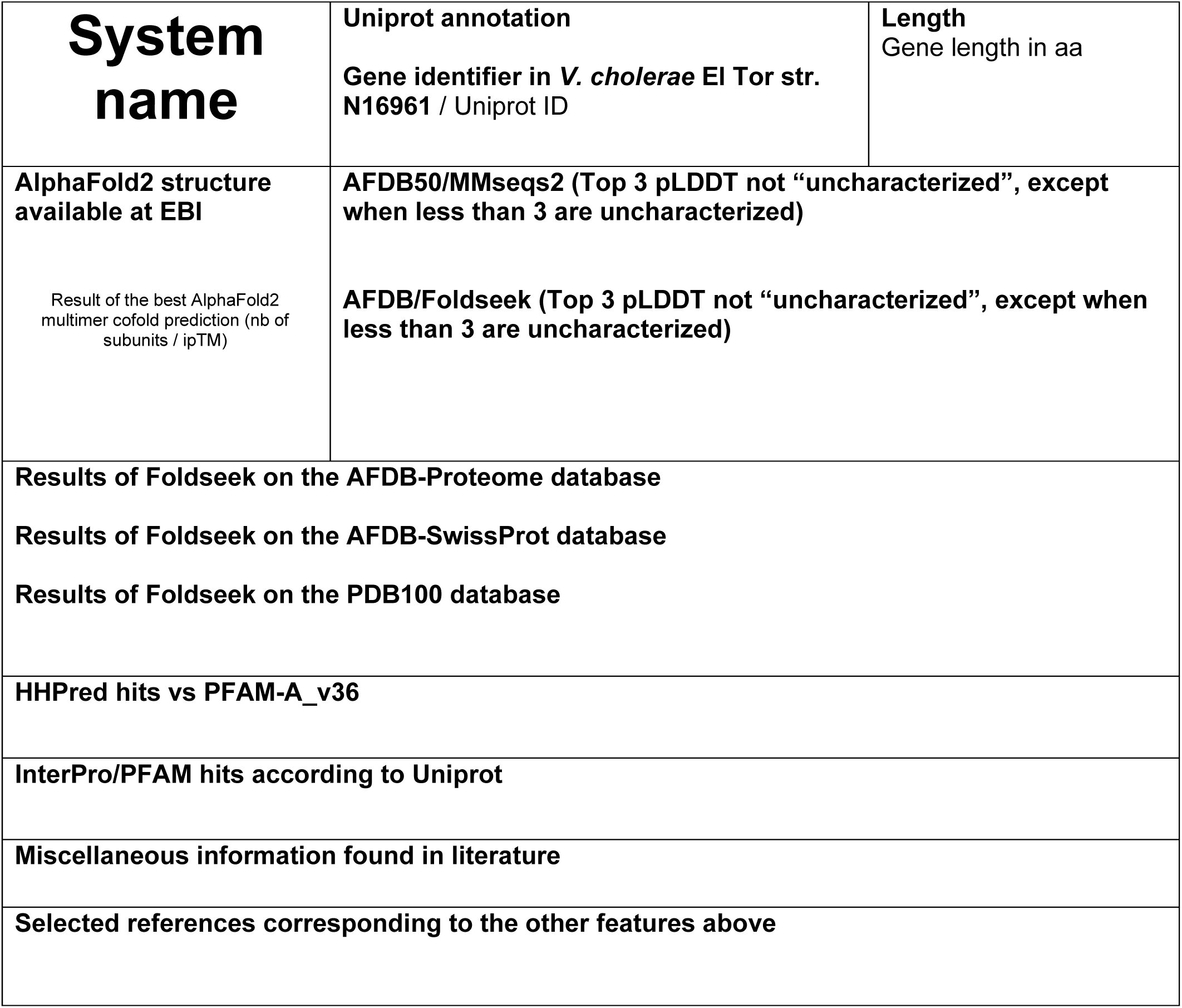
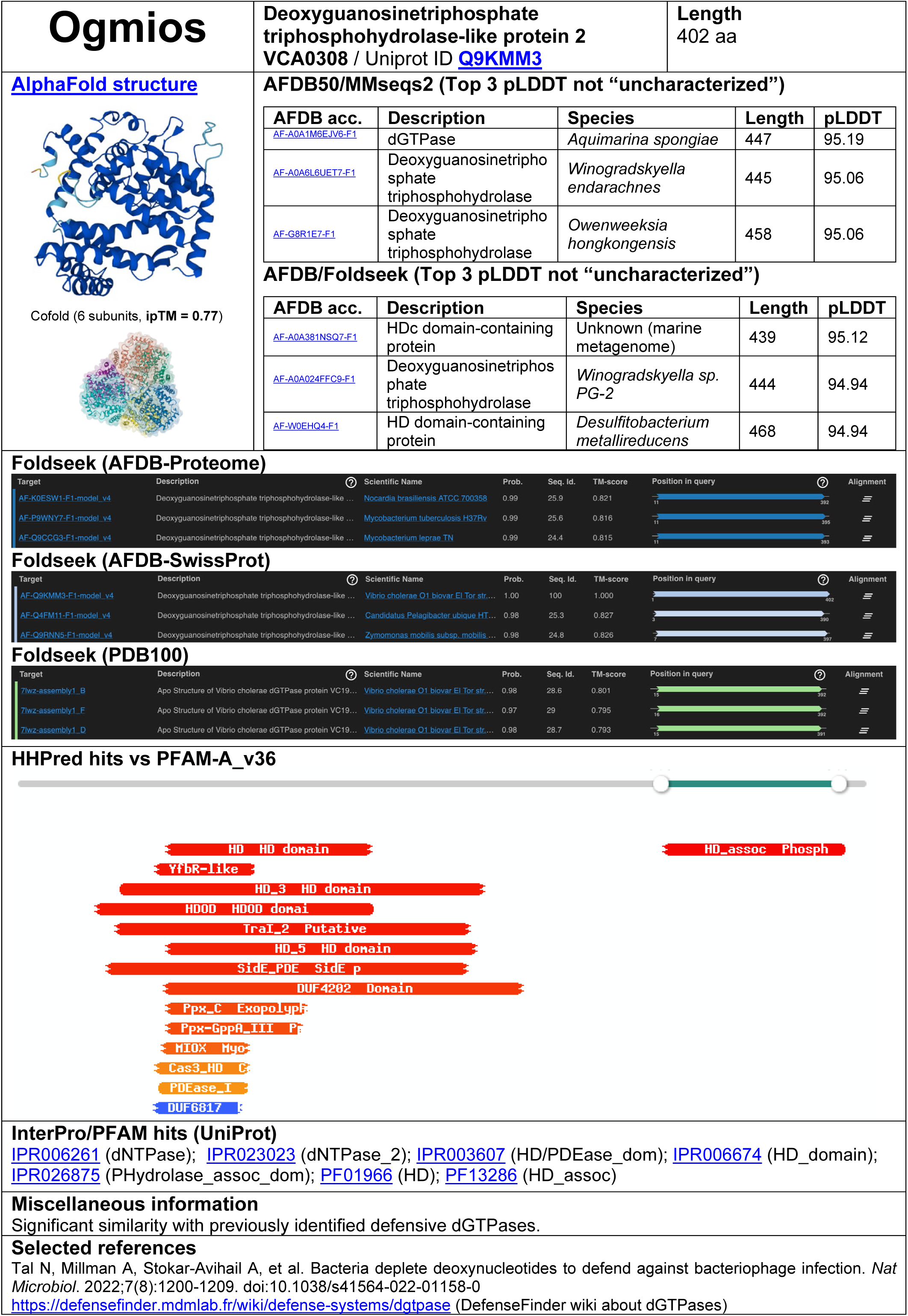
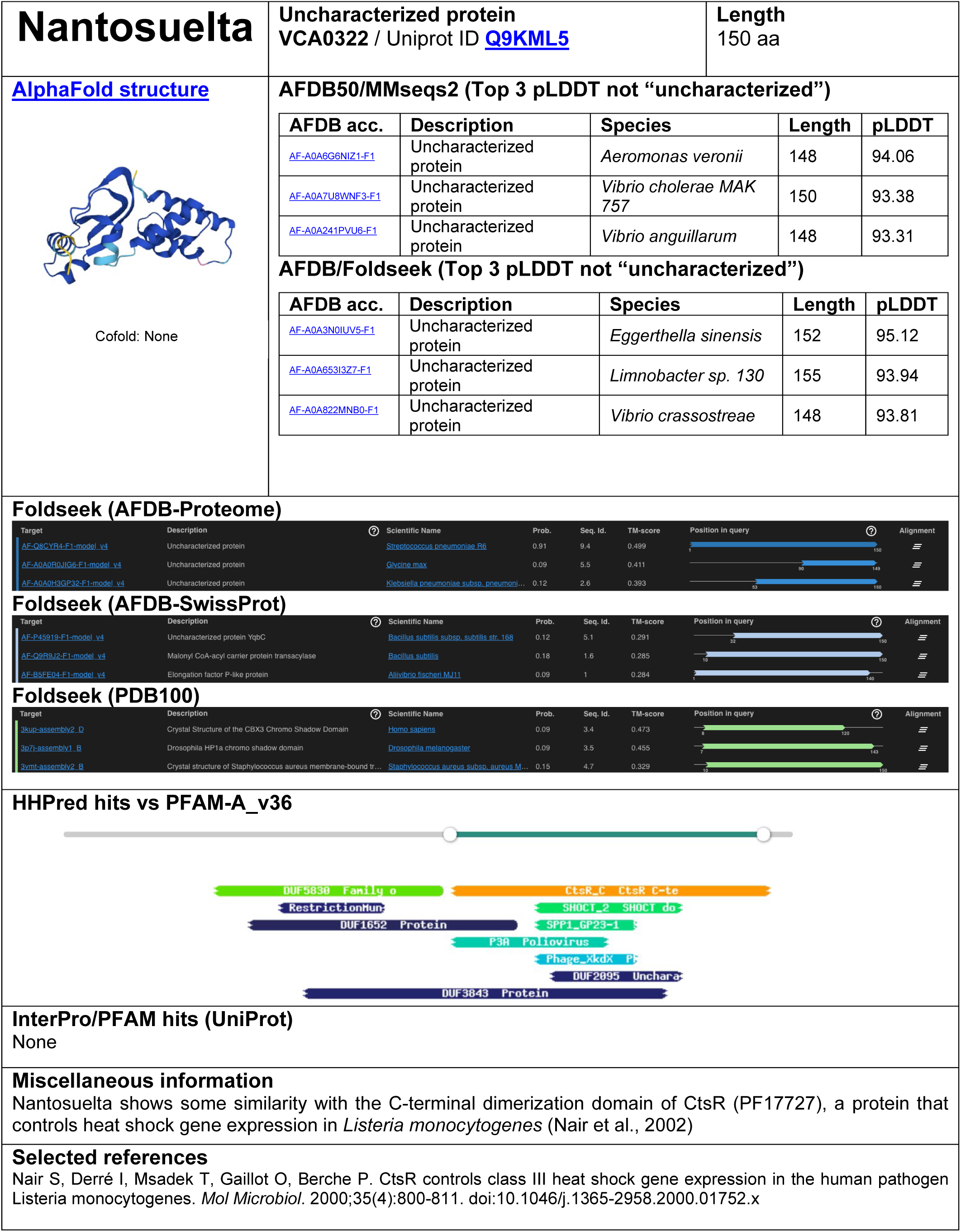
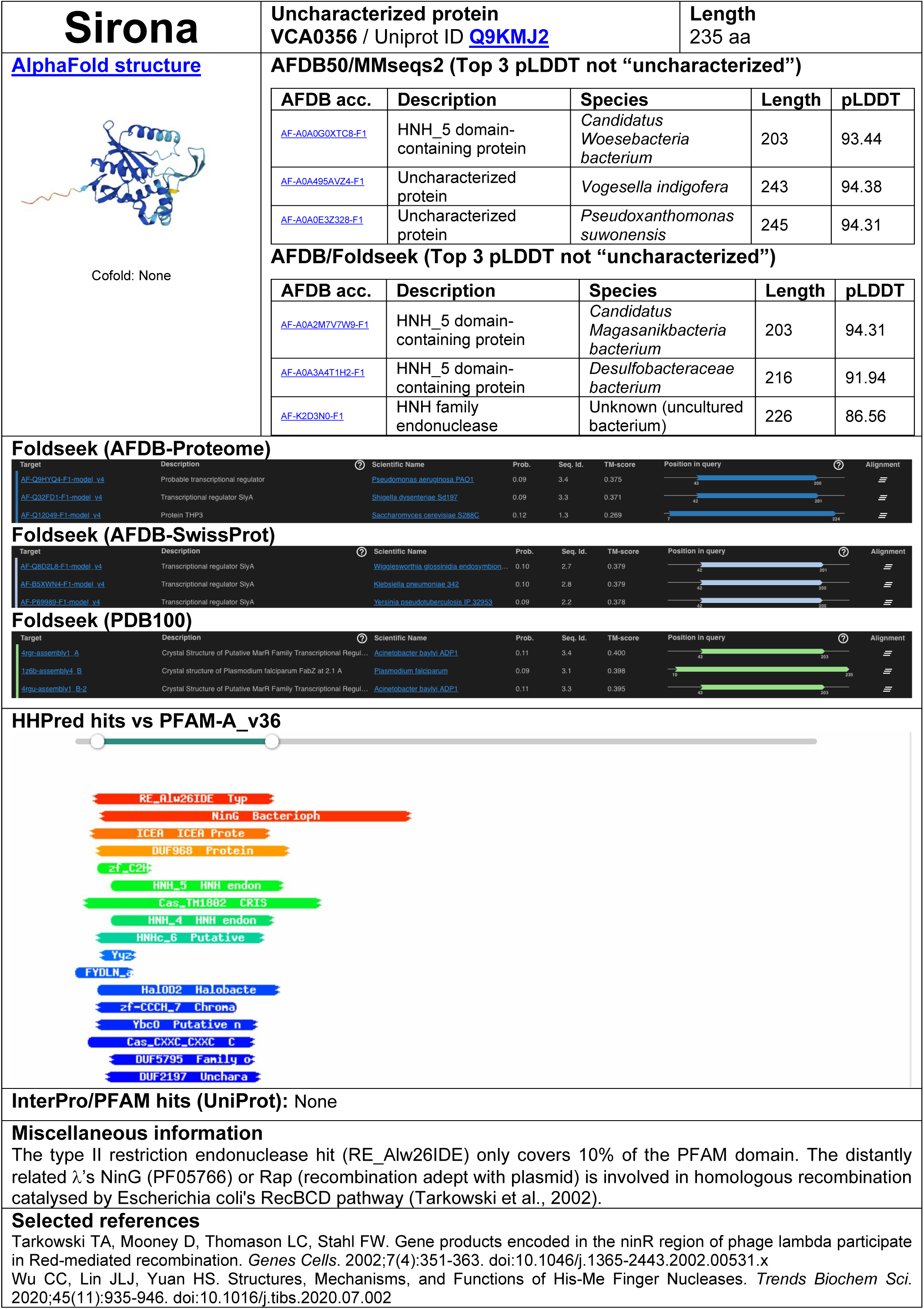
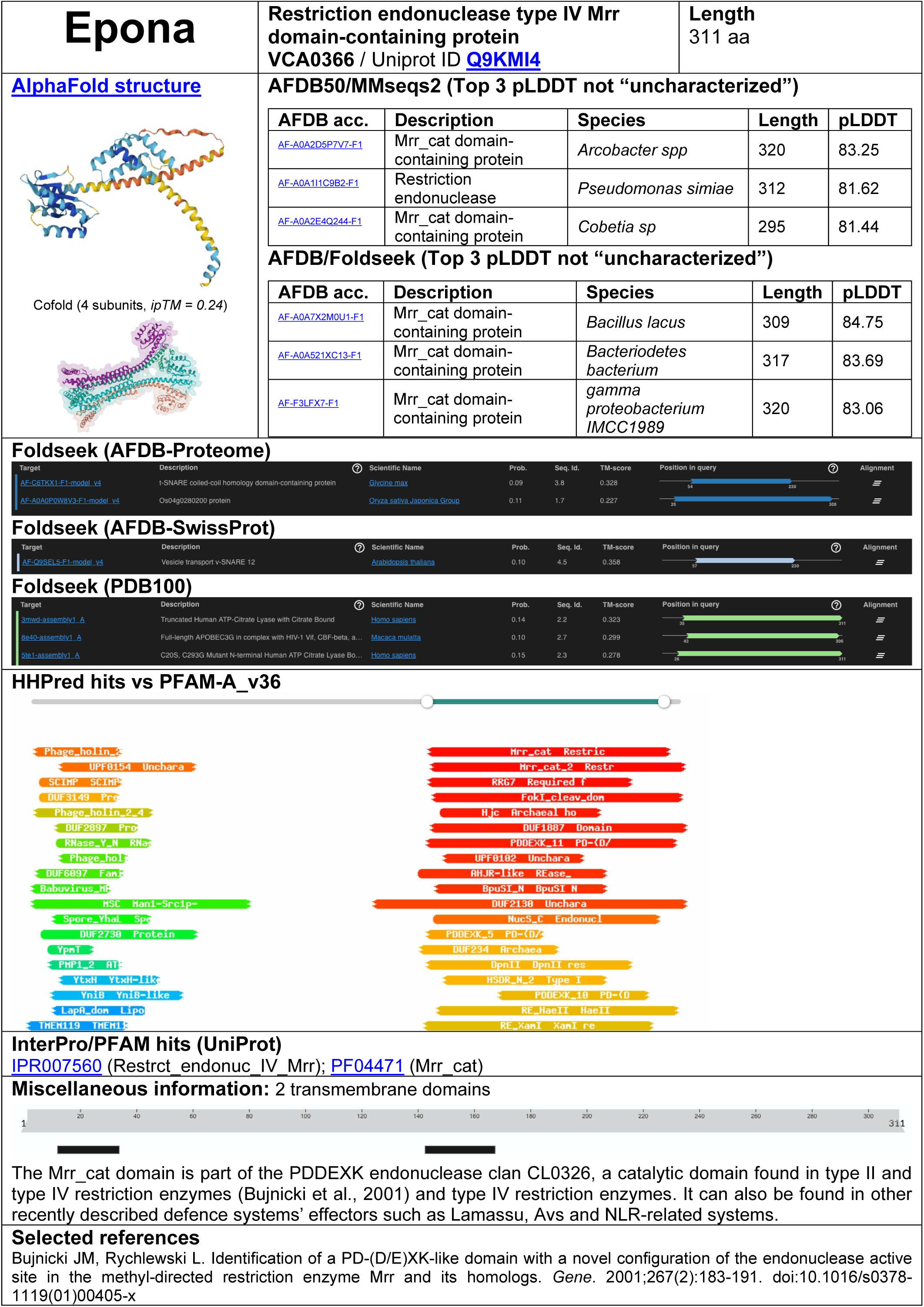
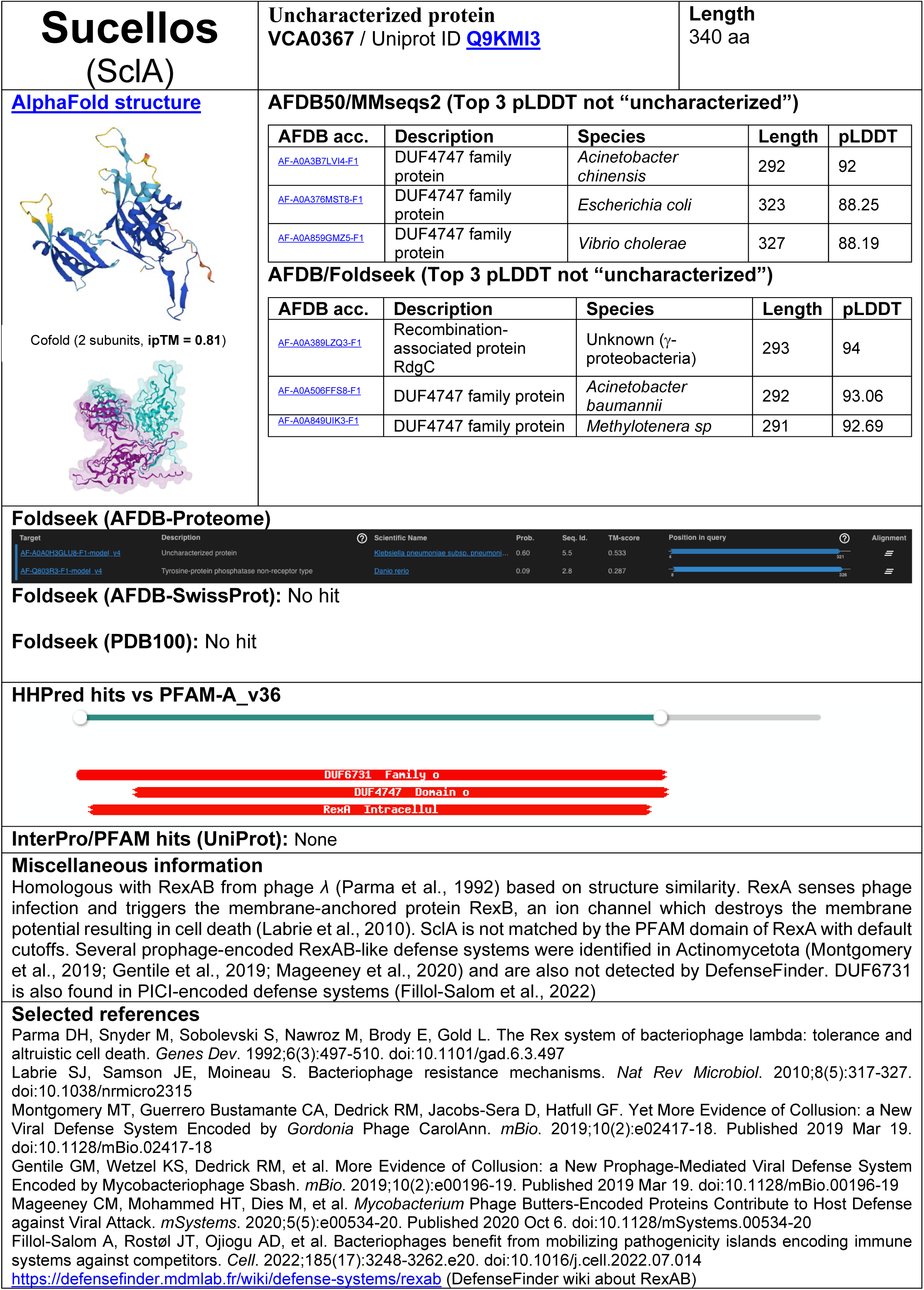
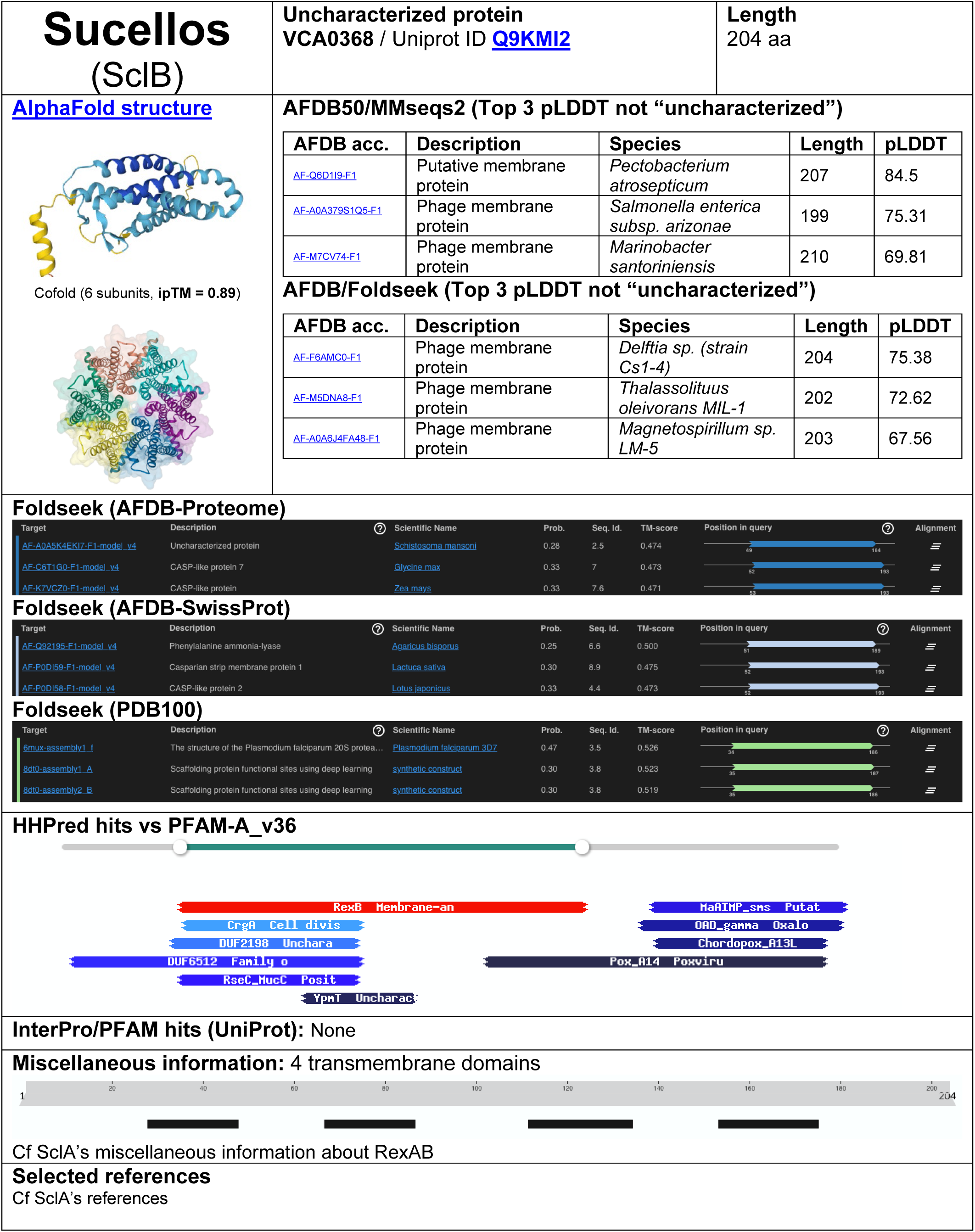
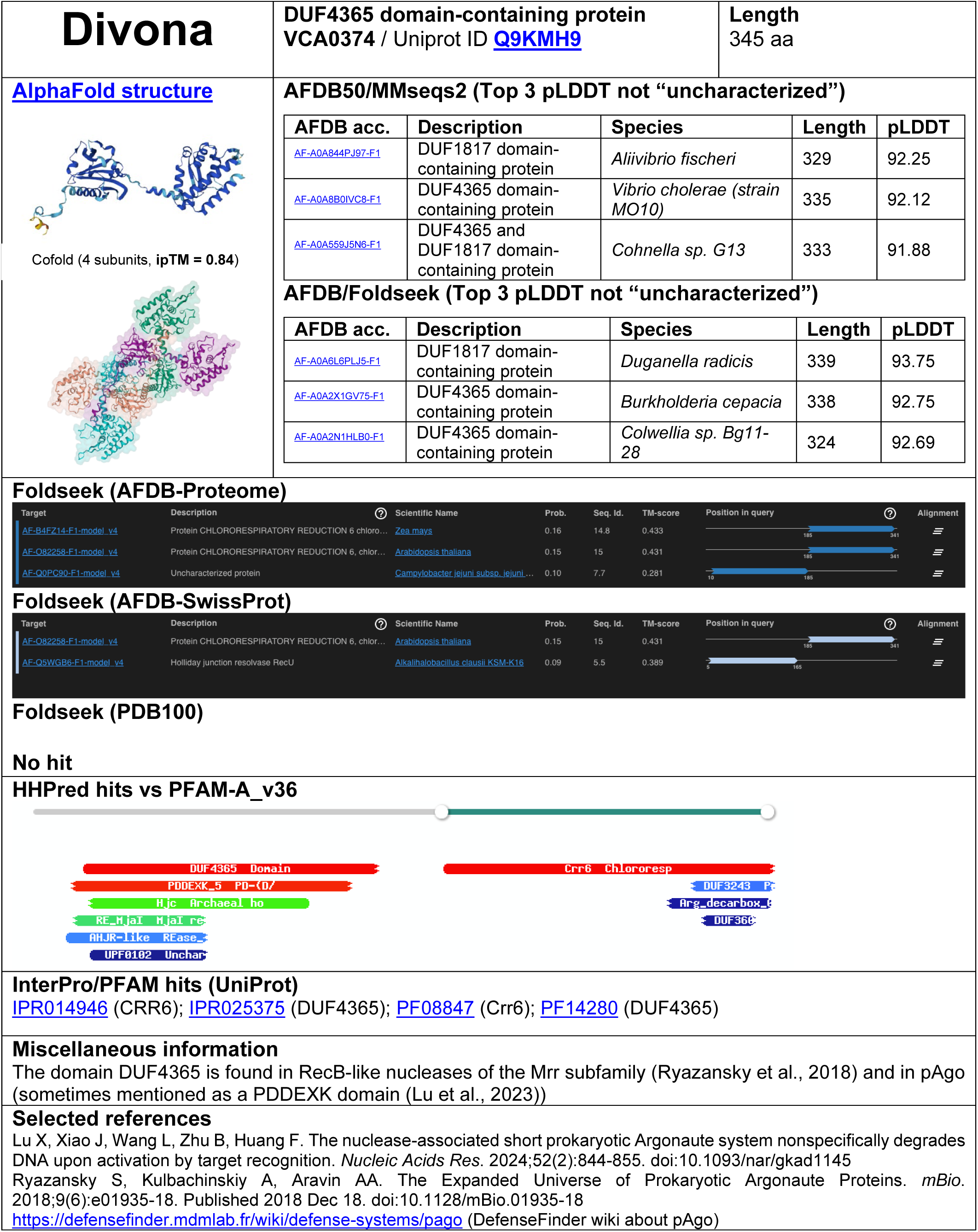
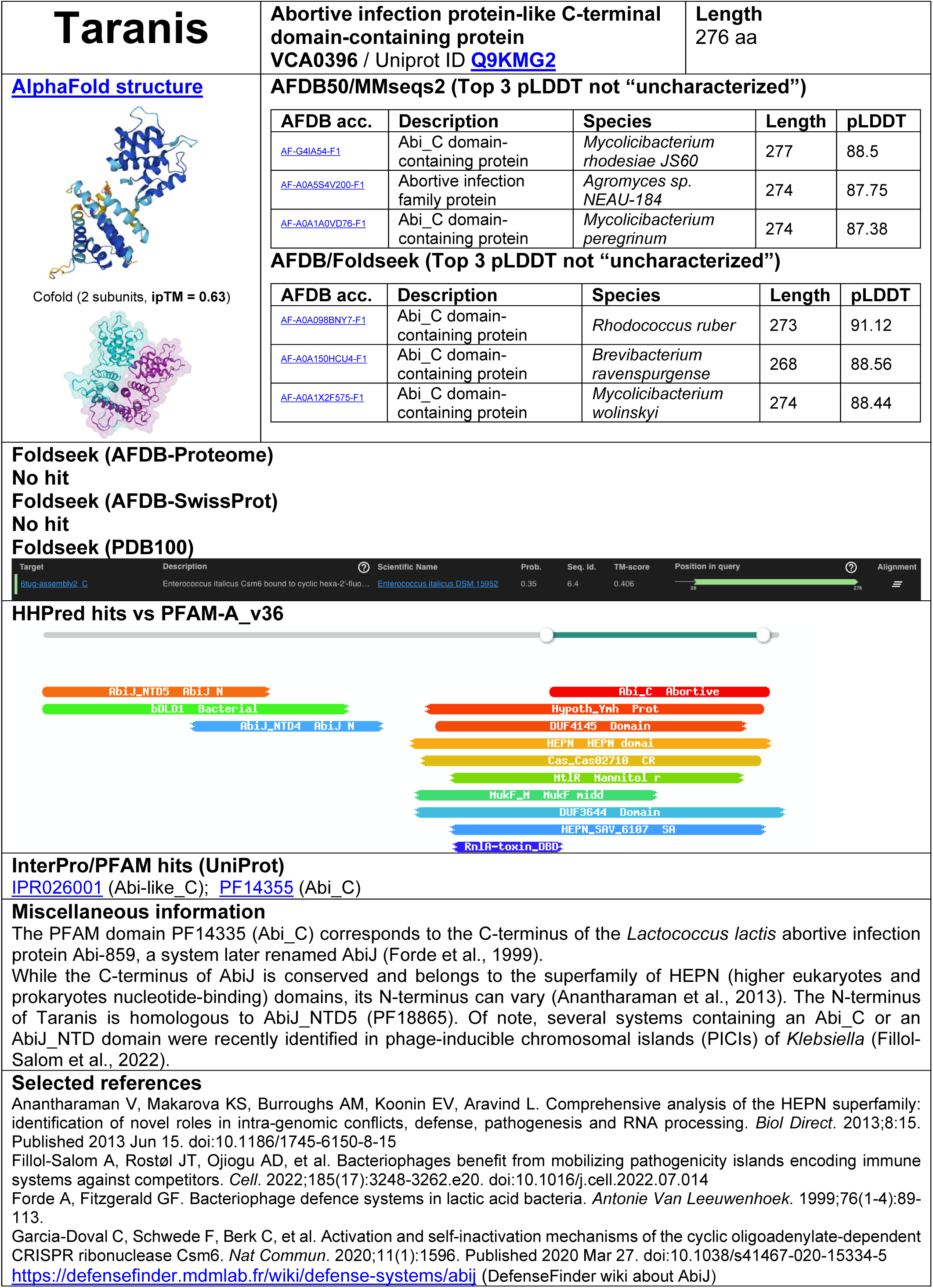
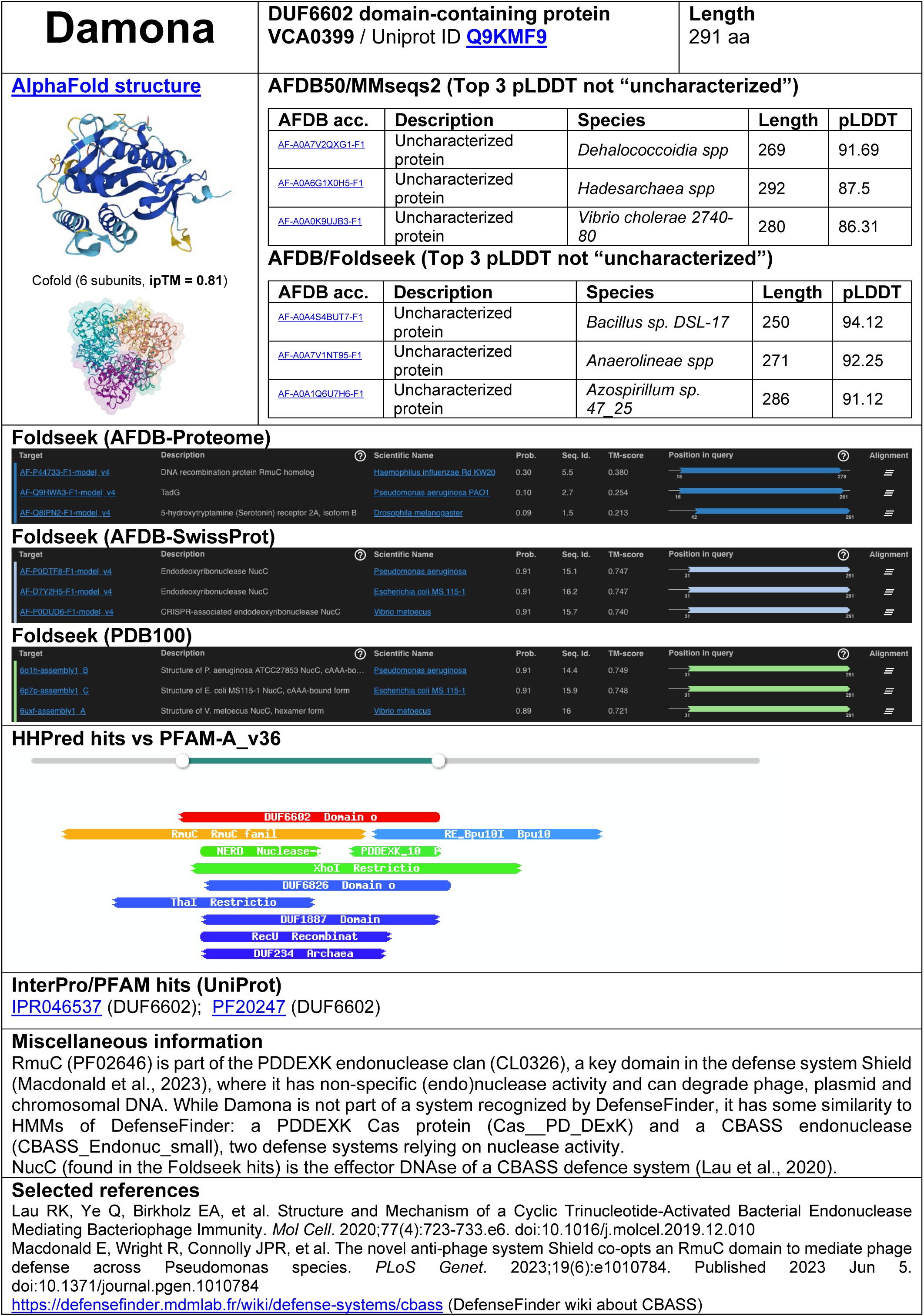
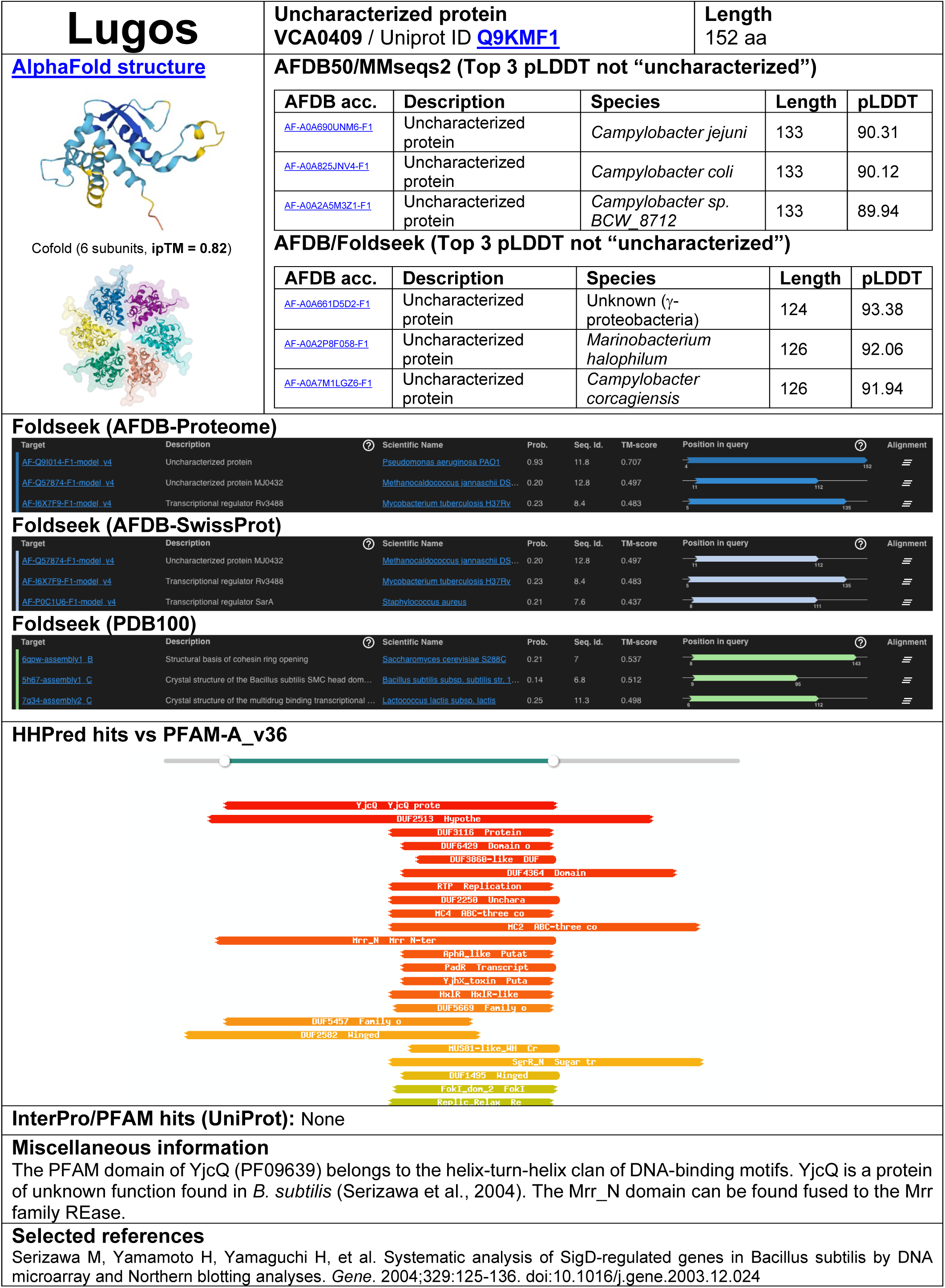
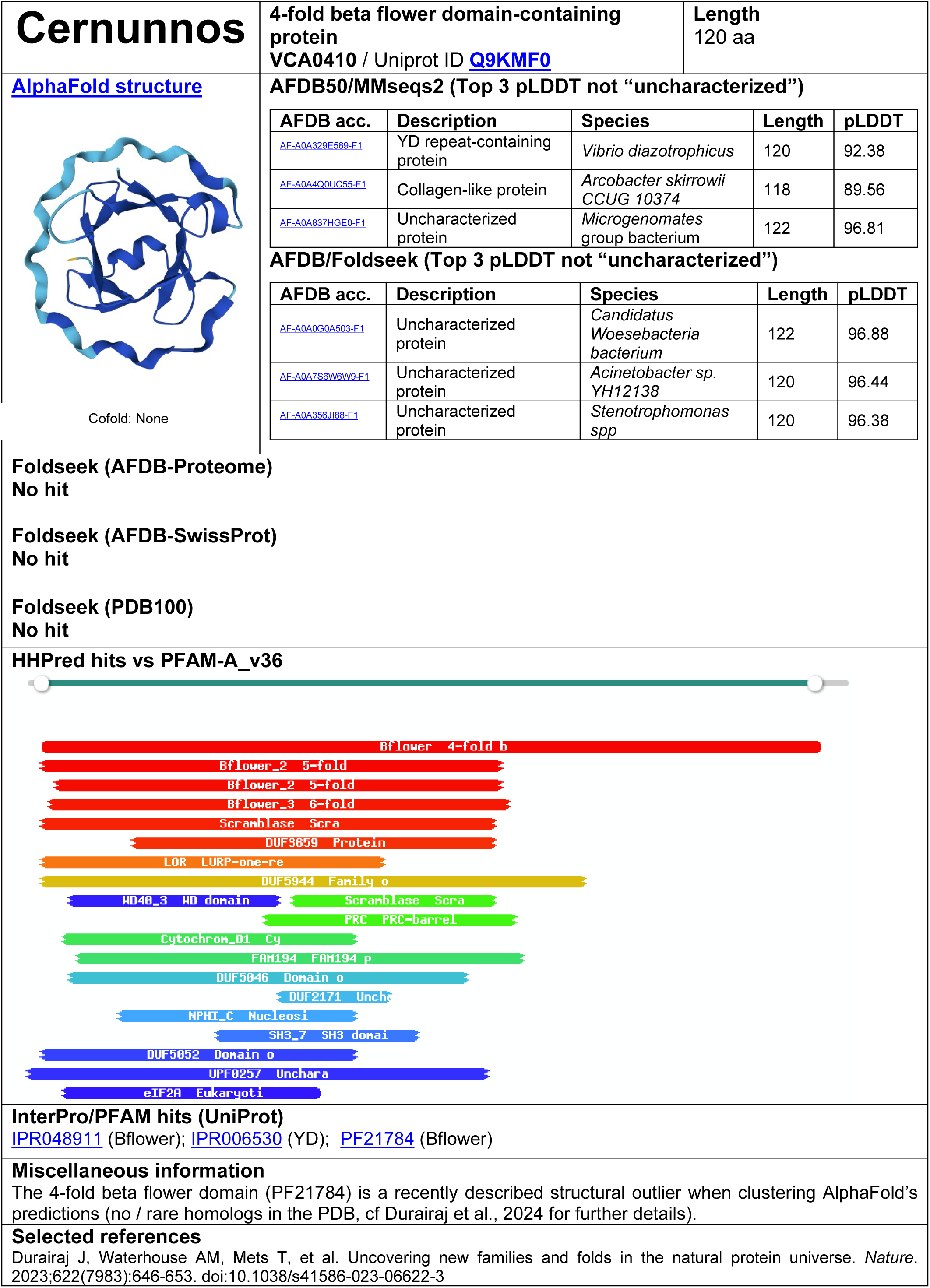
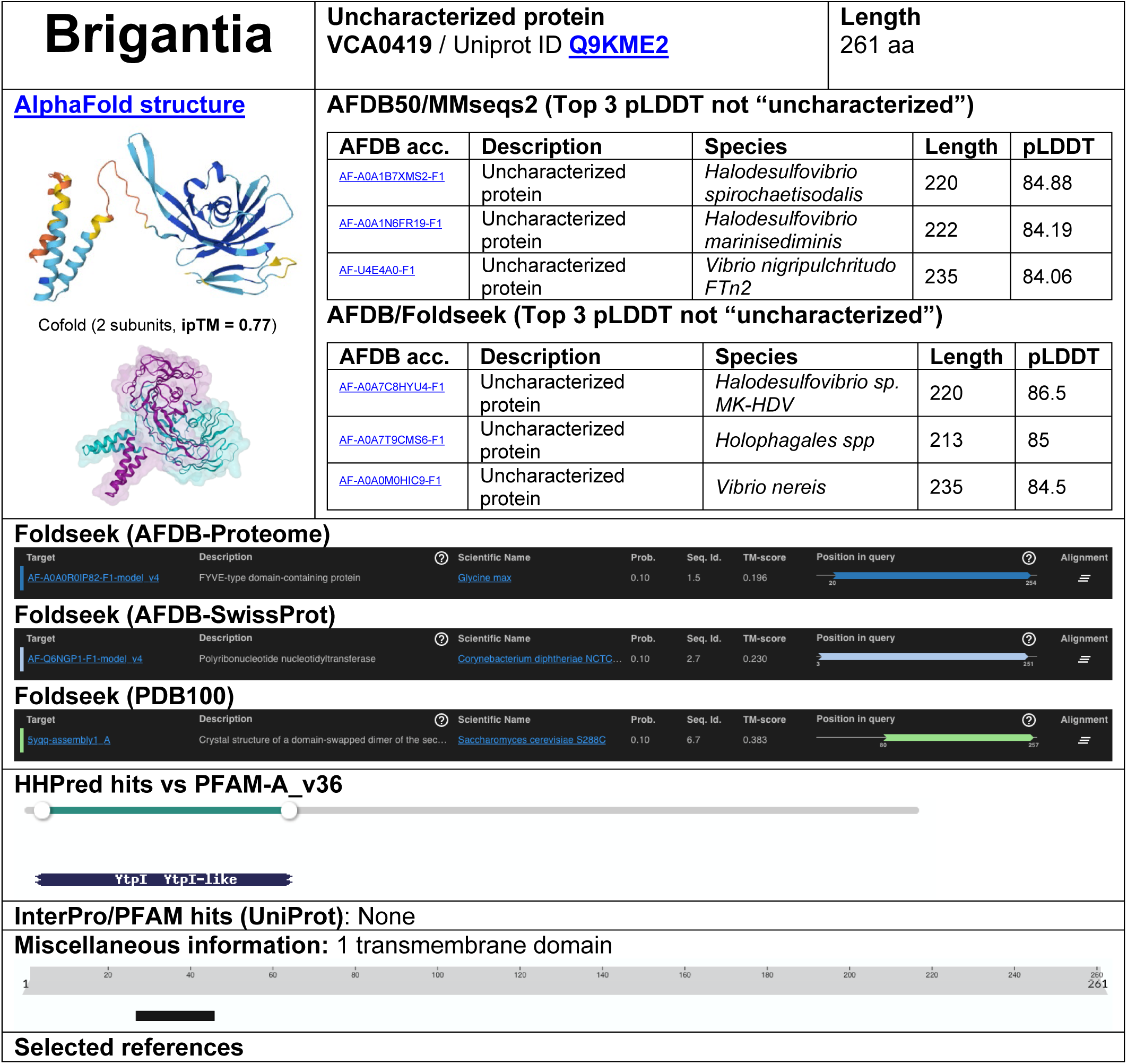
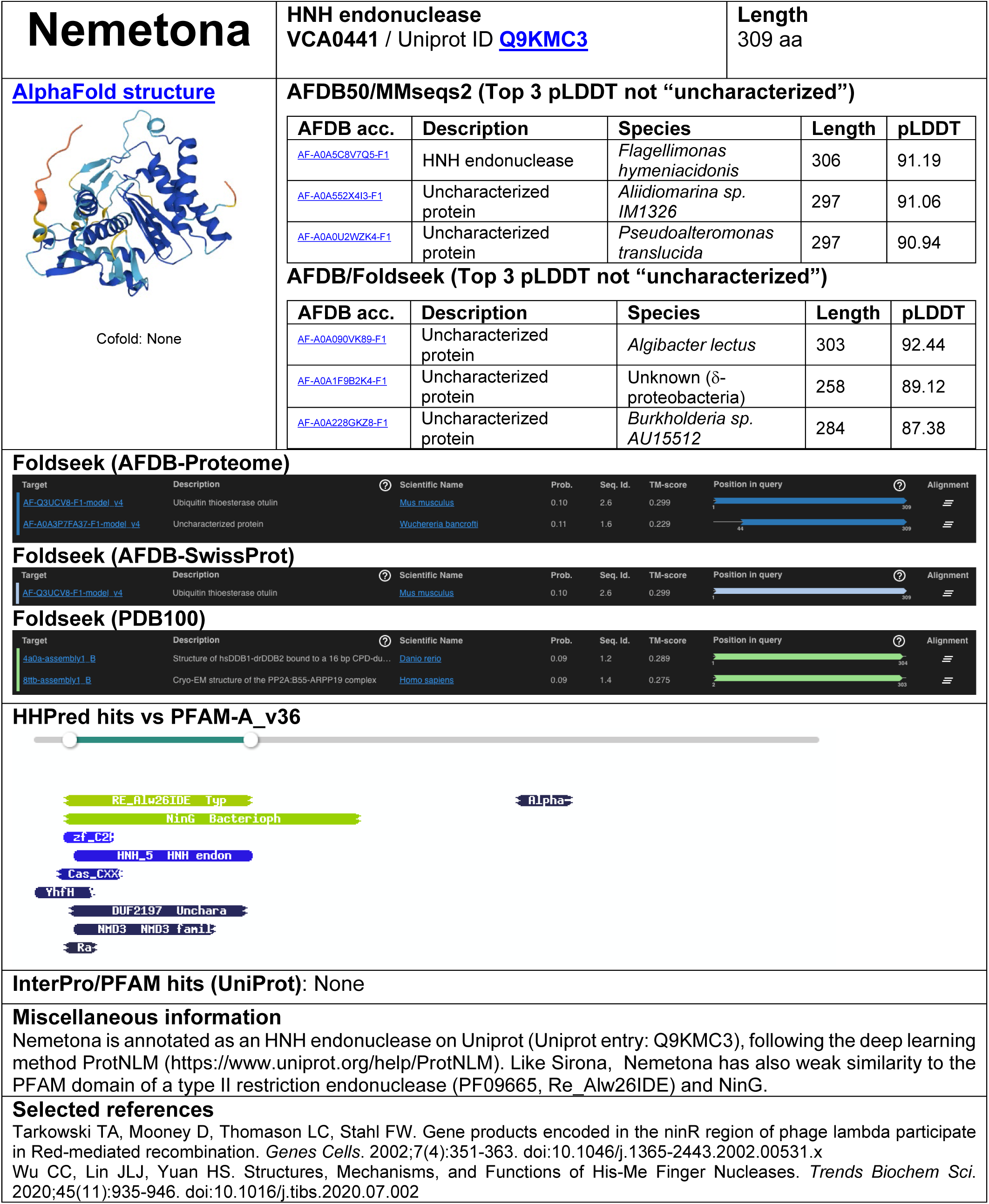
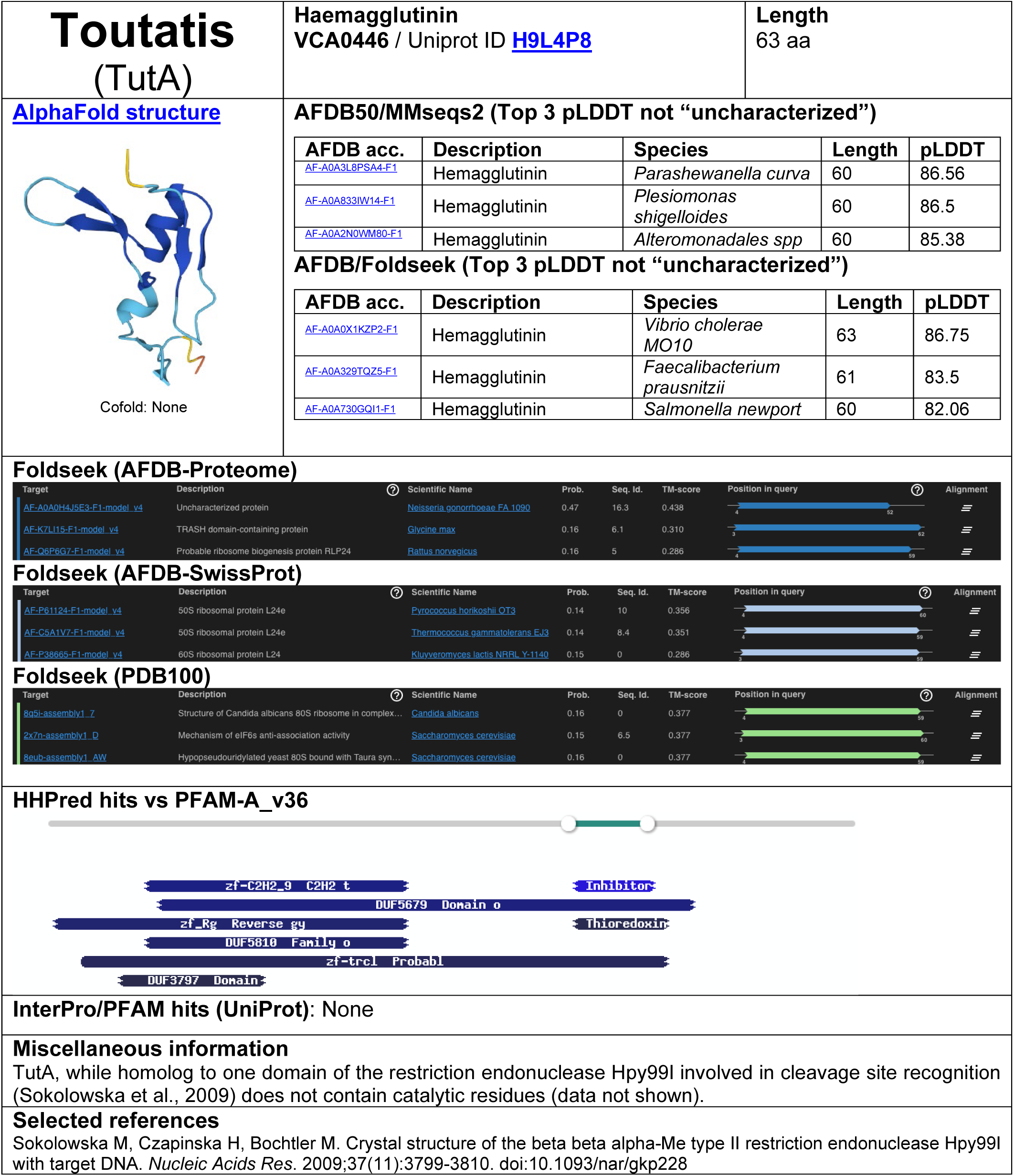
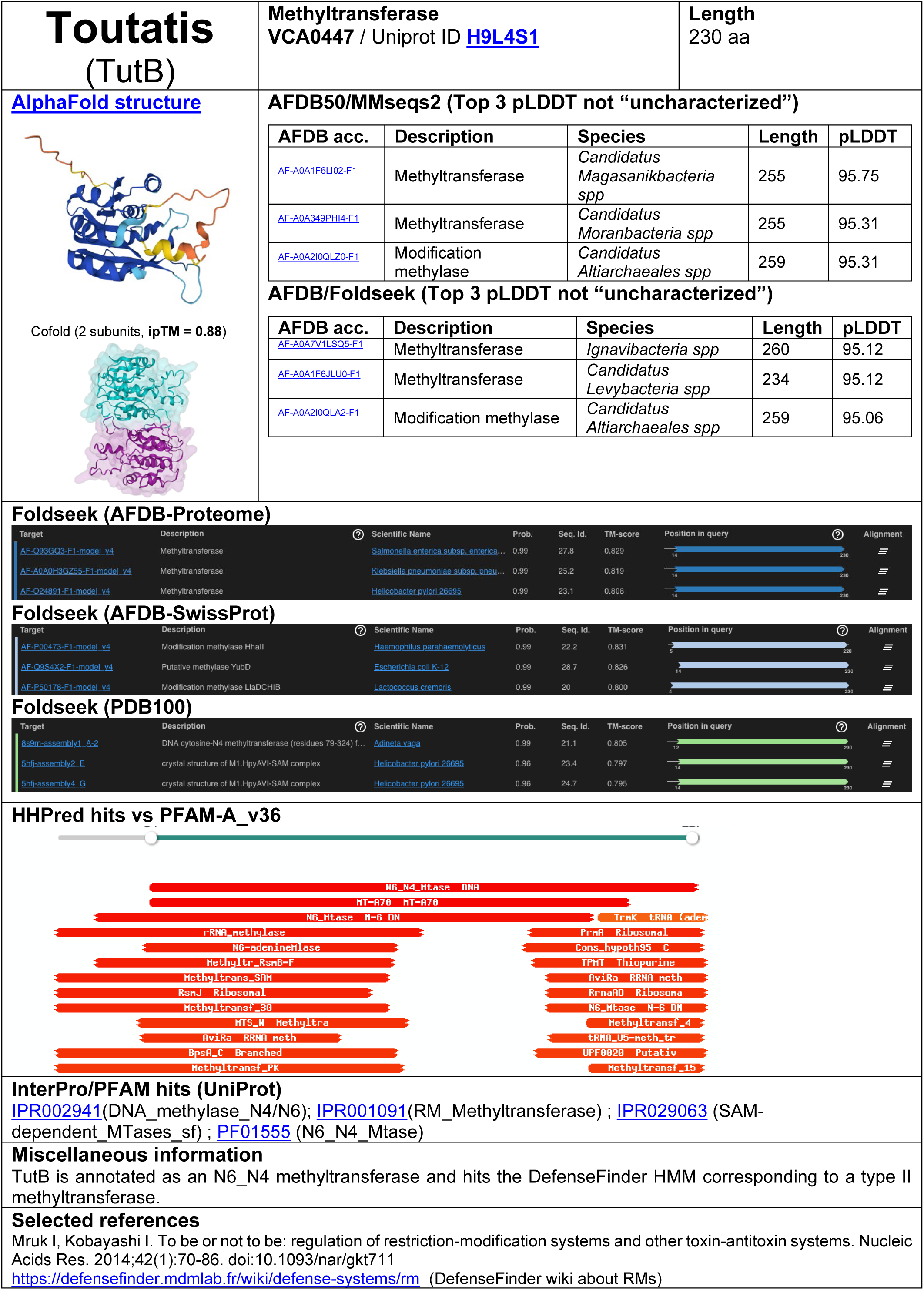
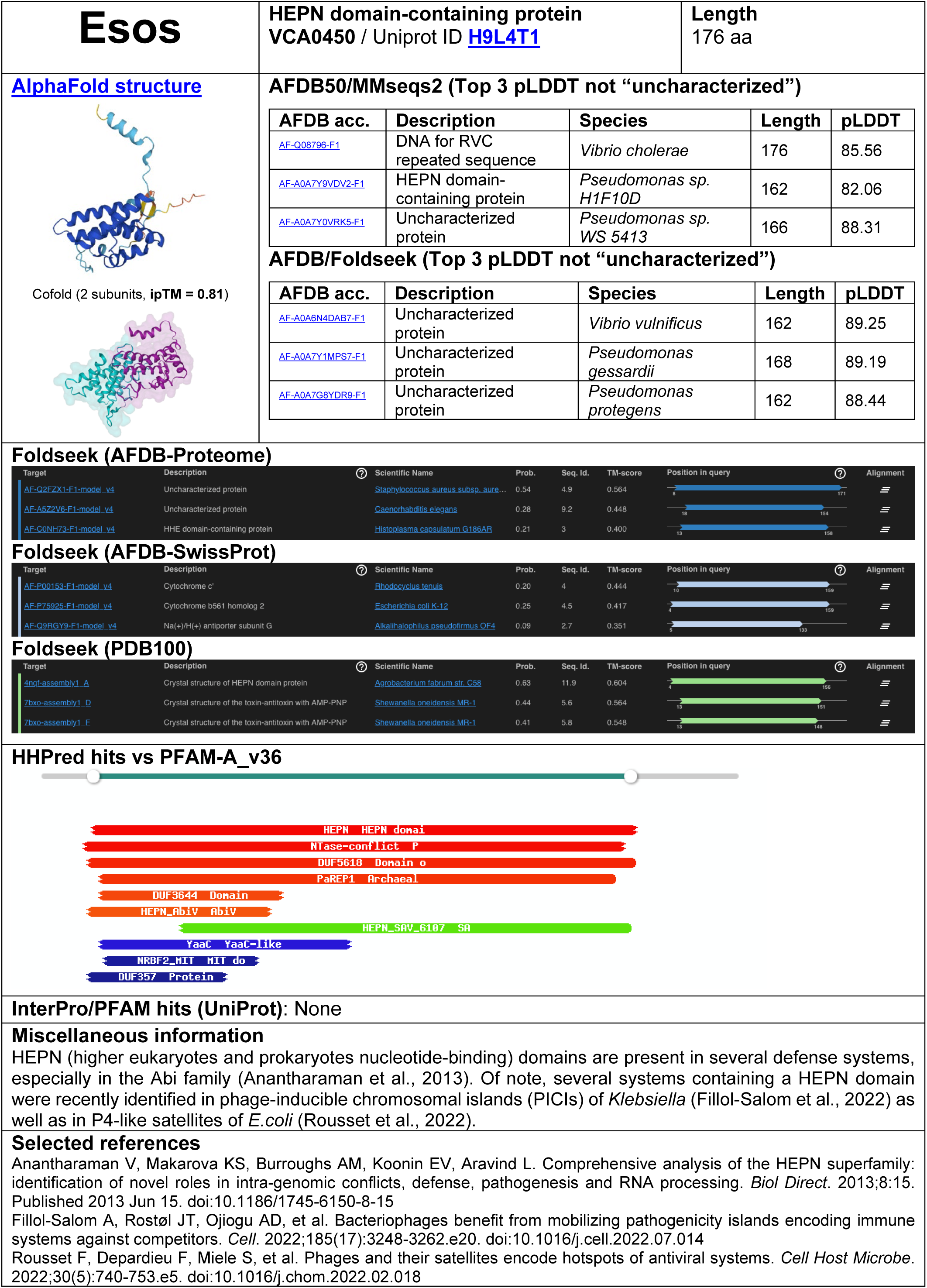
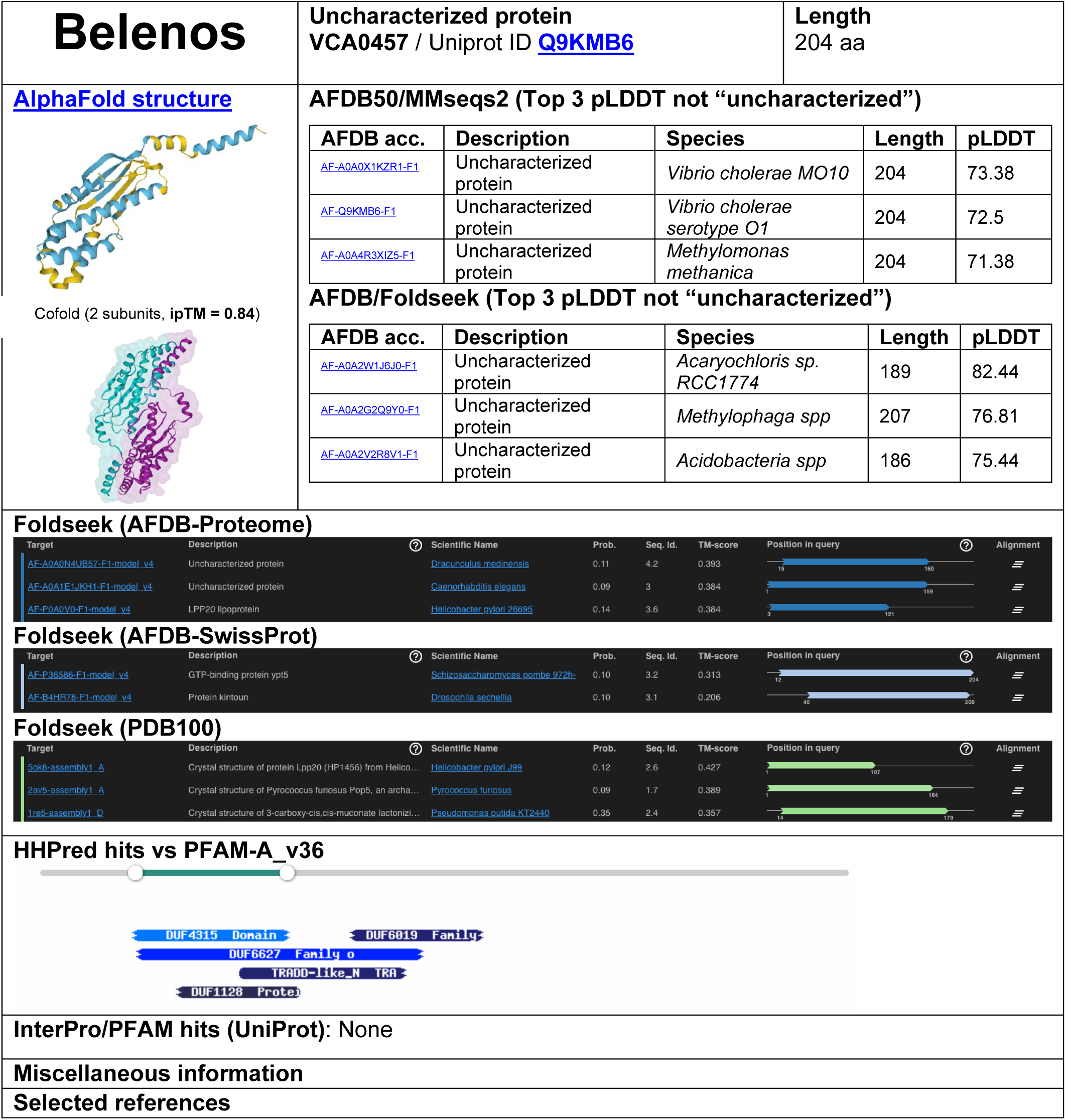
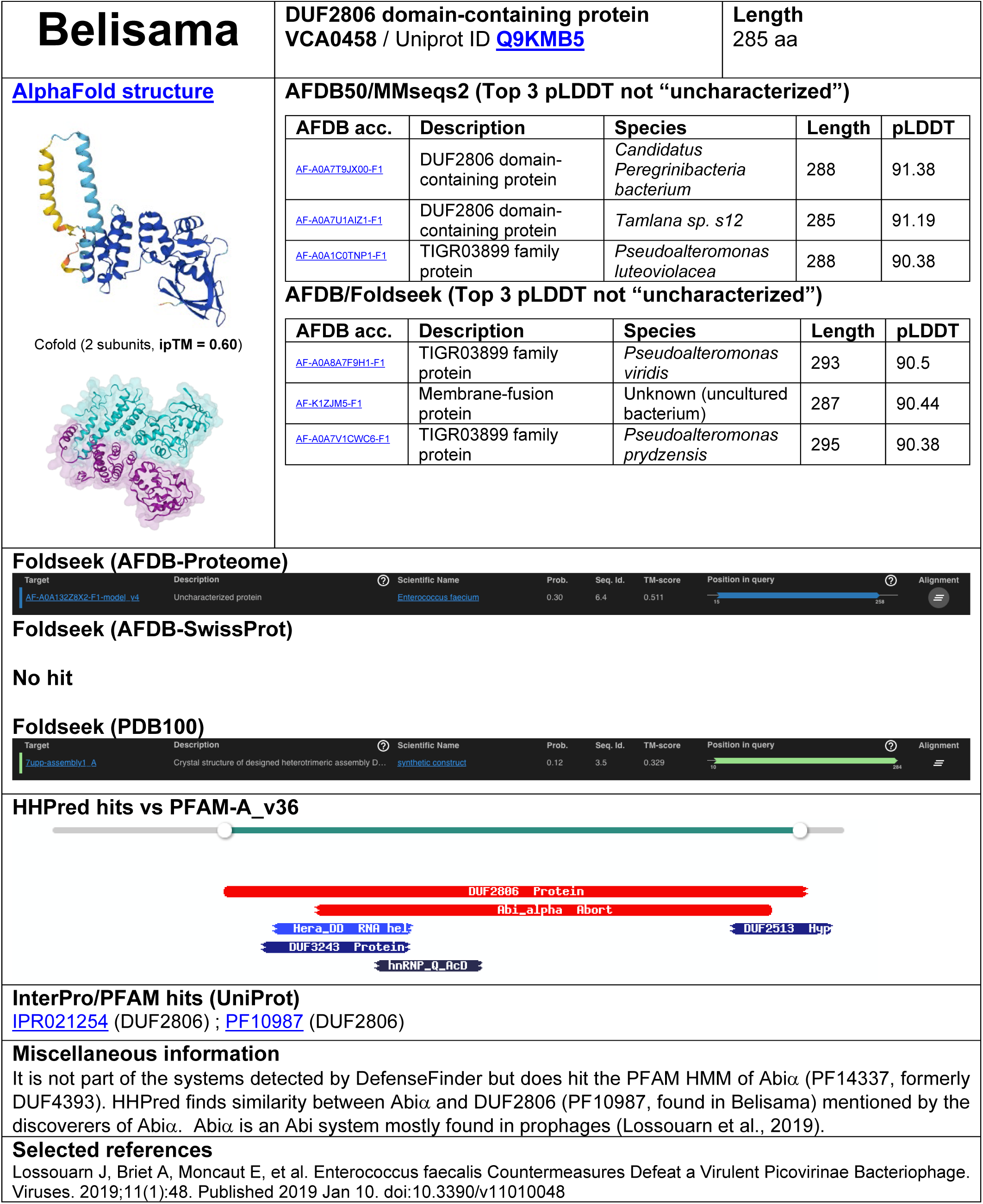
Detailed description of the anti-phage systems found in the *V. cholerae* SCI. All the available data gathered is summarized in an individual sheet by system. The first sheet is a template sheet that explains where each data comes from.

**Figure S5:**
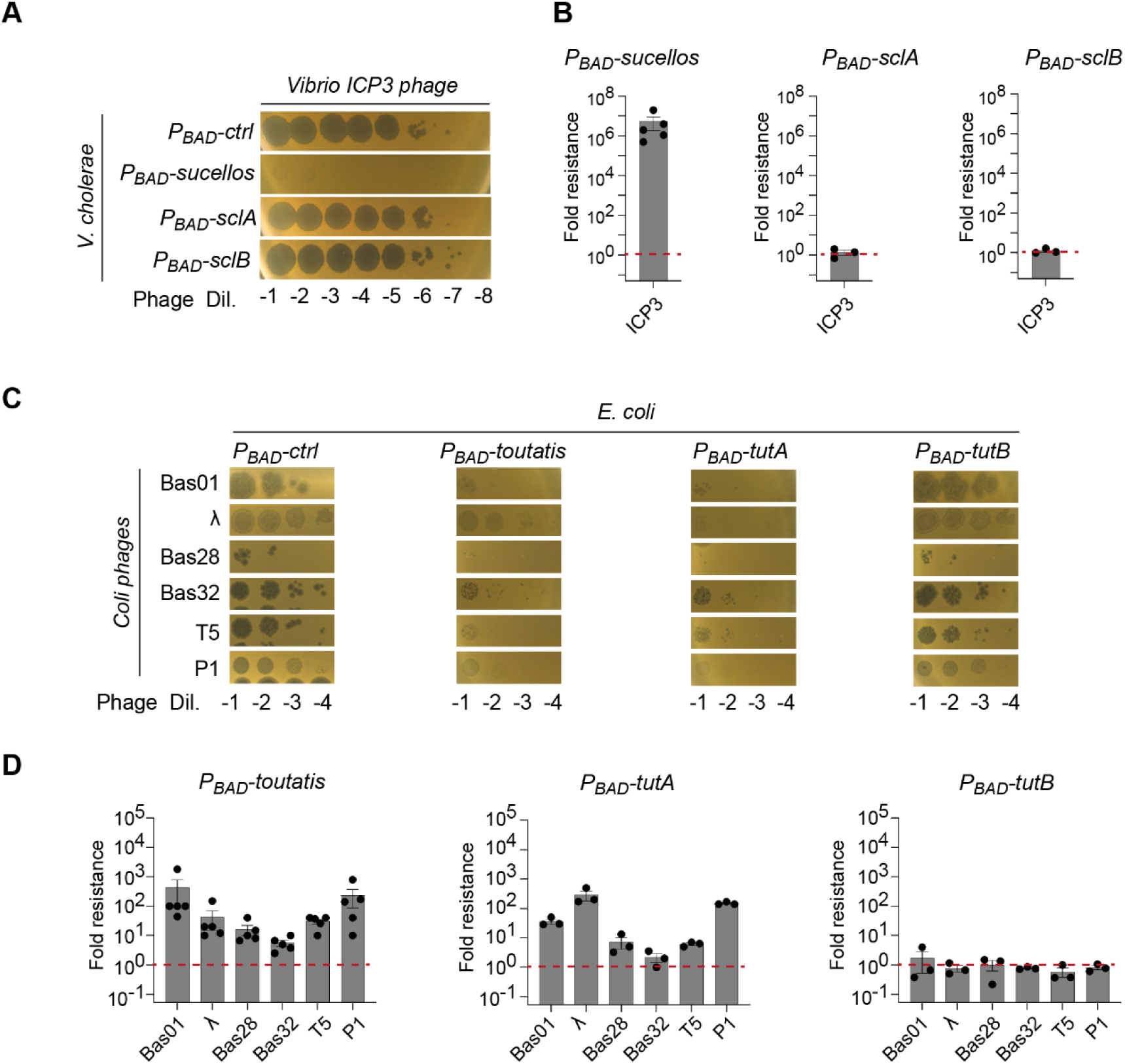
Detailed fold resistance of Sucellos and Toutatis defence systems. **A and C.** 10-fold serial dilutions of a high titer lysate of phages spotted on *V. cholerae* (A) *and E.coli* (C) strains harboring a plasmid expressing the defence systems or not (Редо-cfr/). Representative images from a single replicate out of indepen­dant replicates are shown. **B and D.** Plaque-forming units were measured for each phage on V *cholerae* (B) or *E. coli* (D) strains harboring the control plasmid or the defence systems. Fold-resistance was measured as the ratio between these two values. The bar charts show the mean fold resistance for at least three independent replicates (n>3, individual plots). Error bars shown the standard error of mean (SEM).

**Figure S6:**
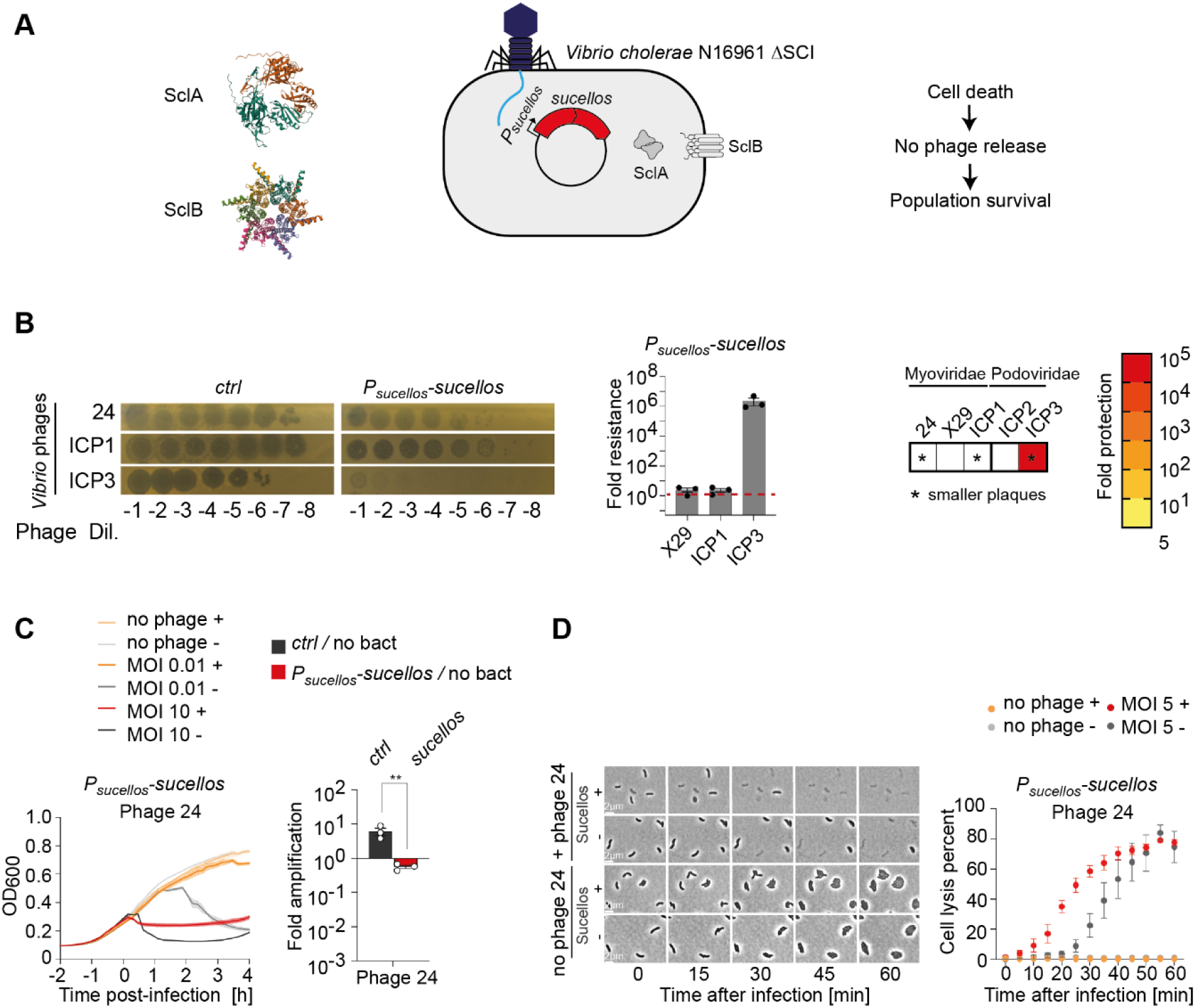
Analysis of the Sucellos system. **A.** AlphaFold structure prediction of the SdA and SclB proteins and model for the defence activity. **B.** 10-fold serial dilutions of a high titer lysate of phages spotted on *V. cholerae* strains. Representative images from a single replicate out of three independant replicates are shown (top panel). Plaque-forming units were measured for each phage on the *V. cholerae* strains harbouring a plasmid expressing *sucellos (P_suce_llos-sucellos)* or not *(Ctrl)*. Fold-resistance was measured as the ratio between these two values. The bar charts show the mean fold resistance for three independent replicates (n=3, individual plots). Error bars shown the standard error of mean (SEM) (bottom right panel). Heatmap of the fold resistance phenotype representing the mean fold resistance of the three independent replicates. Phages are indicated and classed according to morphology. * stands fora marked reduction in plaque size. **C.** As in Figure 4A and B. **D.** Analysis using phase contrast microscopy at 0 min and up to 60 min after infection with phage 24 at an MOI of 5. Cell lysis percent was analyzed every 5 min. Each dot represents the mean of three independent experiments and error bars show the standard error of mean (SEM). The top row shows *V. cholerae* cells containing the anti-phage system and the bottom one without the anti-phage system. Just after phage adding, phage-mediated cell lysis is observed in cells in which the system is present. Representative images from a single replicate out of three independent replicates are shown.

**Figure S7:**
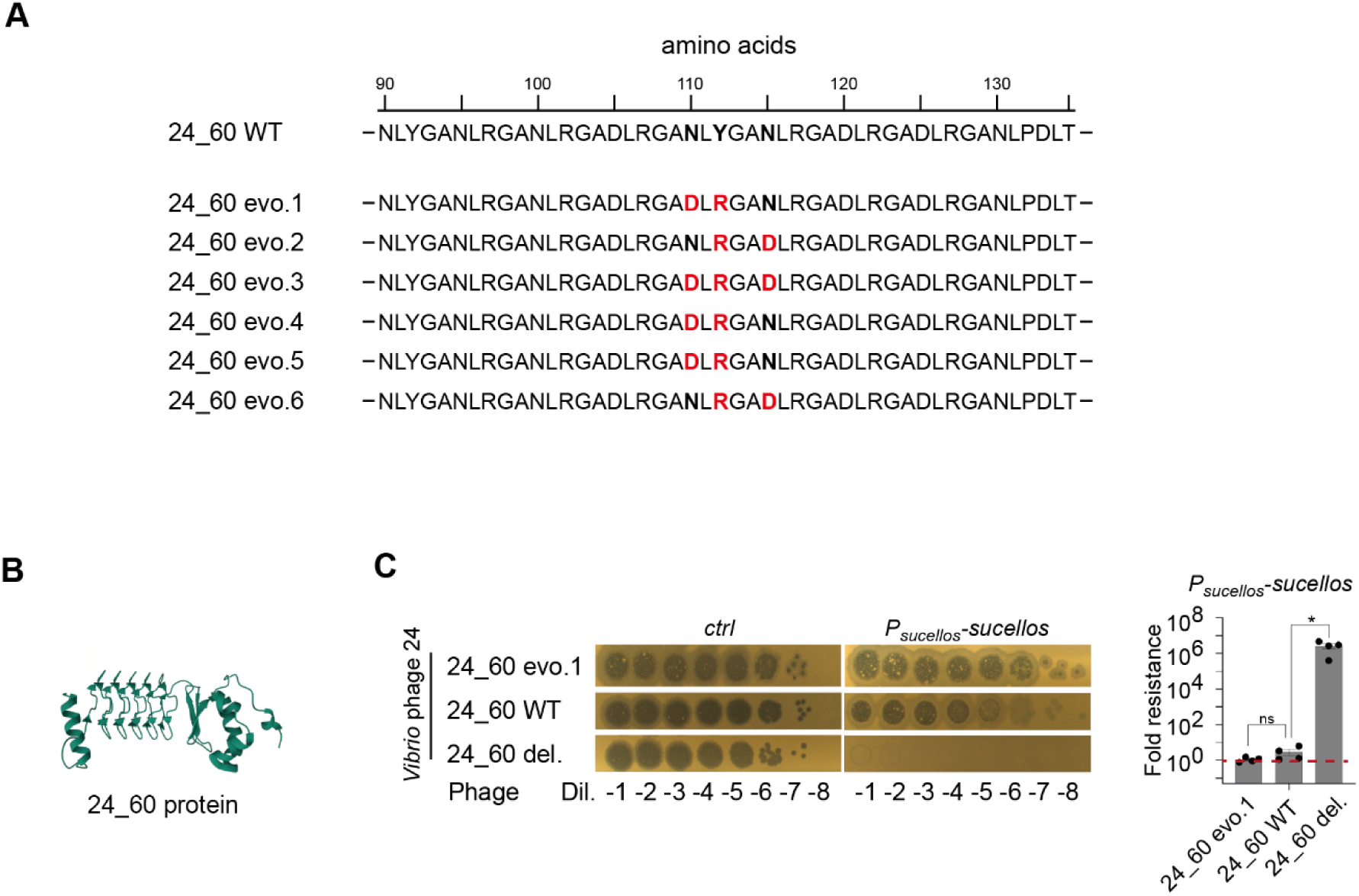
Analysis of the function of the 24_60 phage protein. **A.** The 24_60 amino acid sequences of the *wt* phage 24 and the 6 escapers phages are shown with mutated residues in red. B. AlphaFold structure prediction of the 24_60 protein **C**. 10-fold serial dilutions of a high titer lysate of phages 24 spotted on *V cholerae* strains. Representative images from a single replicate out of three independant replicates are shown (left panel). Plaque-forming units were measured for each phage 24 on the *V. cholerae* strains harbouring a plasmid expressing *sucellos (P_suce_n_os_-sucellos)* or not *{P_S_ucelios’^ctr^h-* Fold-resistance was measured as the ratio between these two values. The bar charts show the mean fold resistance for four independent replicates (n=4, individual plots). Error bars shown the standard error of mean (SEM) (right panel). Statistical comparisons (Student’s t-test, two-tailed) are as follow: ns, not significant; ‘Pvalue < 0.05.

**Figure S8:**
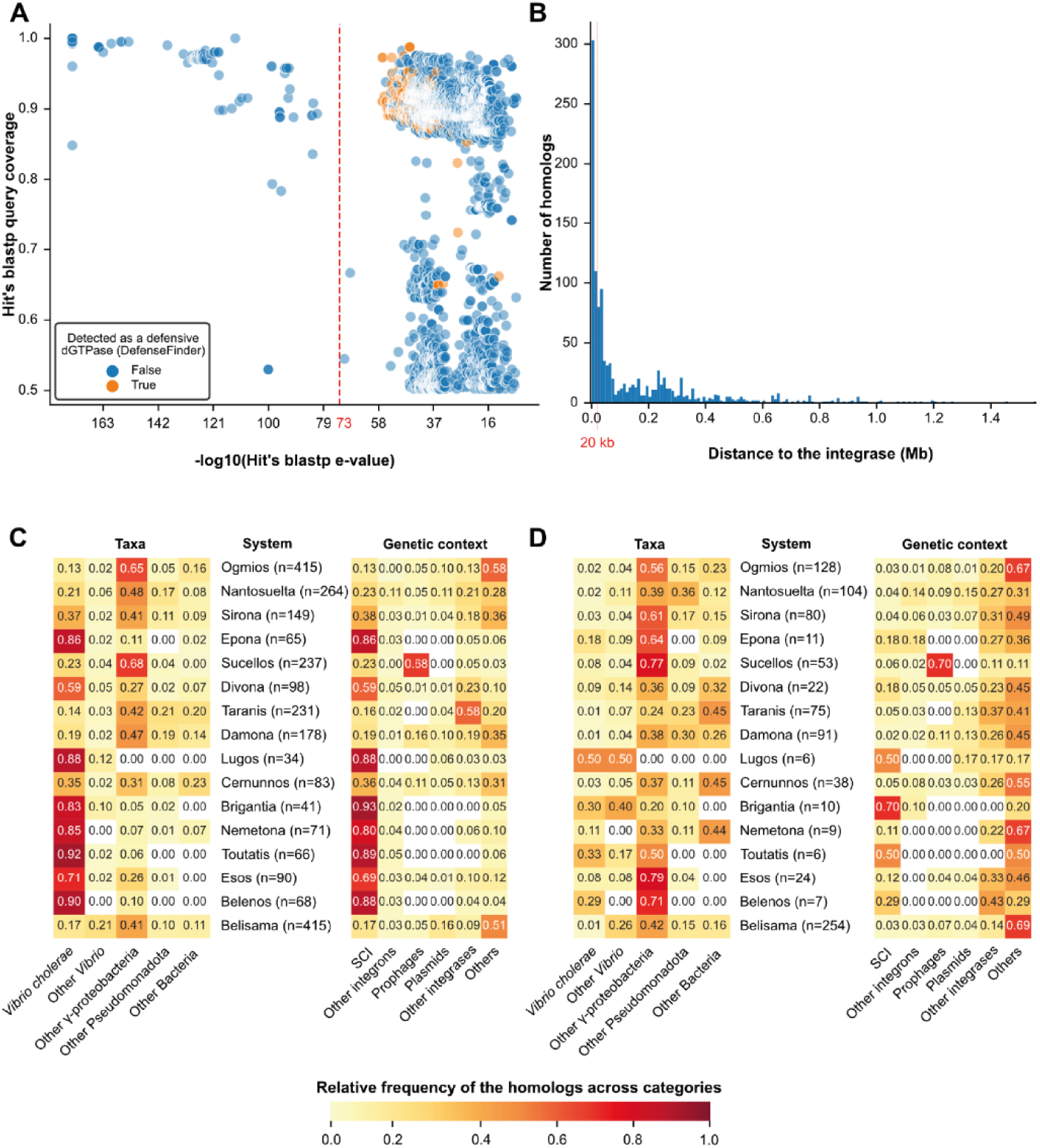
Detailed analysis of the frequency and locations of the newly discovered anti-phage defence systems. **A.** Scatter plot of the homologs of Ogmios identified with blastp. All the hits with a query coverage greater than 0.5 and an e-value smaller than 10-5 (initial thresholds) are displayed. The homologs classified as a defensive dGTPase by DefenseFinder are indicated in orange. The e-value threshold specifically selected for Ogmios homologs is indicated as a red vertical line (10–73). **B.** Histogram of the smallest distance of a system homolog to an integrase. The distance has been computed for the 1175 homologs that were not found to be located in integrons, prophages nor plasmids. A vertical red line indicates a ‘distance of less than 20 kb, retained as the threshold for being close to an integrase. **C.** Detailed version of Figure 5B. Heatmaps of the relative frequency of the novel defence systems across taxa (left) and genetic contexts (right). A number indicates the rounded value of the frequency in each square. The scale of colours is given in the bottom of the graph. The number of systems homologs is indicated after the systems’ names. **D.** Similar analysis as C, except that the homologs were clustered with an identity threshold of 95% (coverage threshold of 80%) to remove redundancy. The number of unique systems homologs (at these thresholds) is indicated after the systems’ names.

